# Spanve: A Statistical Method for Detecting Downstream-Friendly Spatially Variable Genes in Large-Scale Spatial Transcriptomic Data

**DOI:** 10.1101/2023.02.08.527623

**Authors:** Guoxin Cai, Yichang Chen, Shuqing Chen, Xun Gu, Zhan Zhou

## Abstract

Depicting gene expression in a spatial context through spatial transcriptomics would be beneficial for inferring cell function mechanisms. The identification of spatially variable genes is a crucial step in leveraging the spatial transcriptome to understand intricate spatial dynamics. In this study, we developed Spanve, a nonparametric statistical method for detecting spatially variable genes in large-scale ST data by quantifying expression differences between spots and their spatial neighbours. This method offers a nonparametric approach to identifying spatial dependencies in gene expression without assuming specific distributions. Compared to traditional methods, Spanve decreases the number of false-positive outcomes, leading to more accurate identification of spatially variable genes. Furthermore, Spanve could facilitate downstream spatial transcriptomics analyses, including spatial domain detection and cell type deconvolution. These results show the broad applications of Spanve in advancing our understanding of spatial gene expression patterns within complex tissue microenvironments.

**Graphical abstract:** 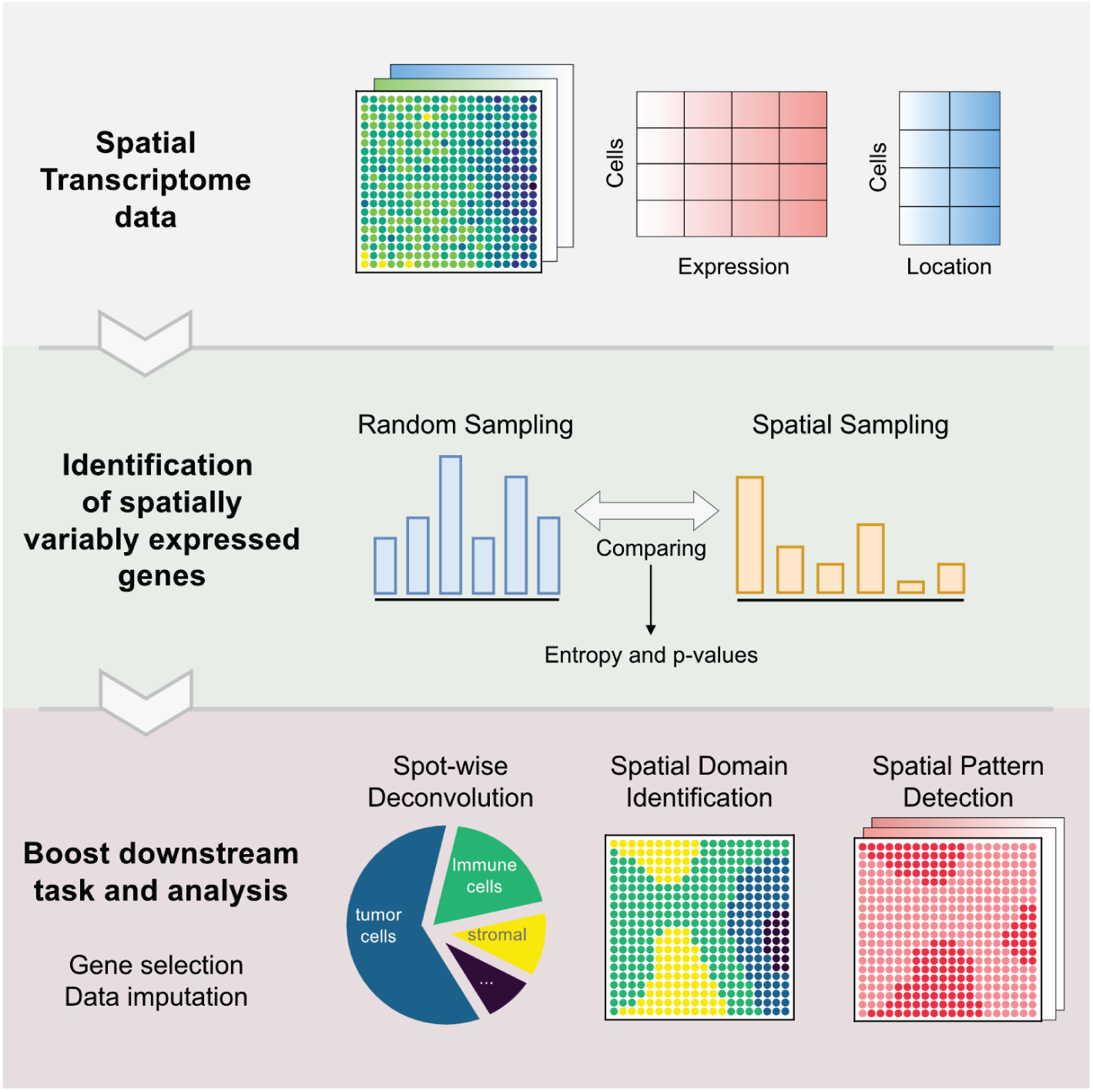

## Introduction

Spatial transcriptomics (ST) maps mRNA molecules within tissue sections to their original locations using arrays of spatially barcoded primers on spots, leading to significant progress in various areas over the past few years ^1,2^. To describe intricate ST landscapes, identification of genes that exhibit differences in expression patterns across different locations within a tissue, which are defined as spatially variable (SV) genes, is crucial. Unlike highly variable genes (HVGs), which may vary between cells or groups of cells, SV genes exhibit spatial patterns across spatial locations. These spatial patterns are often characterized by distinct spatial domains where the expression of SV genes is homogenous, indicating shared functional attributes, cell composition or structural characteristics. Thus, SV genes can reflect a myriad of factors, including the localization of cell types or space-dependent interactions between cells ^1^, providing valuable biological signals for the complex interplay between gene expression, spatial organization, and functional processes within tissues and ultimately advancing our understanding of biological systems and their intricate spatial dynamics.

As SV genes may contain important information about the underlying biological mechanisms within tissues, several studies have attempted to detect SV genes by modeling gene expression. The most common strategy uses Gaussian regression processes that assume gene expression follows a specific probability distribution, such as Gaussian ^3^ or Poisson ^4^, with a covariance matrix composed of spatial and nonspatial components. However, whether gene expression can be well described by these classical distributions is questionable. There is also a similar problem in single-cell RNA sequencing (scRNA-seq) data due to overdispersion and drop-out events ^5^. Two specialized distributions (negative binomial, NB, and zero-inflated negative binomial, ZINB) have been proposed to solve this problem. As these distributions may not adequately consider spatial effects on gene expression, algorithms based on the distribution hypothesis may lead to false-positive outcomes when modeling ST data, especially for genes with high spatial heterogeneity based on our observations. In other words, genes with low *p* values may not exhibit a spatial variation pattern, which may be caused by distribution assumption violations. Other methods, such as cluster-based ^6,7^ or graph-based methods ^8–10^, avoid arbitrary assumptions. However, these methods may have limitations in handling large-scale data. New sequencing technologies, such as stereo-seq ^11^, allow for the identification of cellular or subcellular transcripts ^12^ but also present new challenges due to the increased volume and complexity of data. Spatial autocorrelation methods, including Moran’s I and Geary’s C ^13,14^, are scalable but have a large false-positive rate in our following experiments and a previous benchmark ^15^.

To this end, we developed SPAtial Neighborhood Variable Expressed gene detection (Spanve), a statistical method for analyzing the dependence of gene expression on spatial location. Spanve was developed to enhance the specificity of SV gene identification without assuming specific distributions while maintaining low computational costs and promoting downstream tasks, such as clustering and deconvolution. Based on previous studies, we established benchmarks for SV gene identification, computational cost and downstream analyses. The results indicate that our proposed method enhances the specificity of identification with low computational costs and promotes downstream tasks, such as clustering and deconvolution. To demonstrate the practical application of Spanve, we analyzed a human breast cancer ST dataset to characterize the intratumor heterogeneity.

Within the Spanve framework, we implemented a spotwise colocalization analysis, which may reveal potential cell‒cell interactions. Furthermore, we explored the associations between SV genes and potential contributing factors to enhance our understanding of underlying biological processes. These findings highlight the ability of Spanve to effectively identify important SV genes and facilitate downstream analyses of ST data.

### Design

We developed Spanve, a non-parametric statistical method for measuring spatial variance in gene expression within ST data. Spanve enhances downstream analyses by reducing false positives in spatially variable gene detection. The fundamental premise of this study is that gene expression differences between spots are homogeneous in the absence of spatial effects. Rather than modeling gene expression directly using classical distributions, Spanve models spatial variance by quantifying differences between a spot and its spatial neighbors. We implemented two approaches for identifying spatial neighbors: k-nearest neighbor-based (Spanve-k) and Delaunay network-based (Spanve-d) methods. By adjusting the number of neighbors, researchers can tailor the resolution of the analysis to their specific aims. The Delaunay-based method is particularly suited for large-scale data analysis.

The selection of spots and their spatial neighbors can be viewed as a form of sampling or network construction based on spatial location. In contrast, the null model randomly selects spot pairs. Genes exhibiting spatial expression patterns will show significant differences between space-based and random sampling. This approach differs from current statistical methods that often compare to assumed distributions, such as Poisson. We posit that sampling from the data itself inherently accounts for background effects, potentially reducing false positives. Moreover, our analysis demonstrates that assumed distributions may not accurately represent gene expression, particularly for SV genes. We quantify the Kullback-Leibler (KL) divergence between the two sampling distributions and perform a statistical test to identify genes with significant spatial variability (details in STAR Methods, Figure 1A).

**Figure 1.**
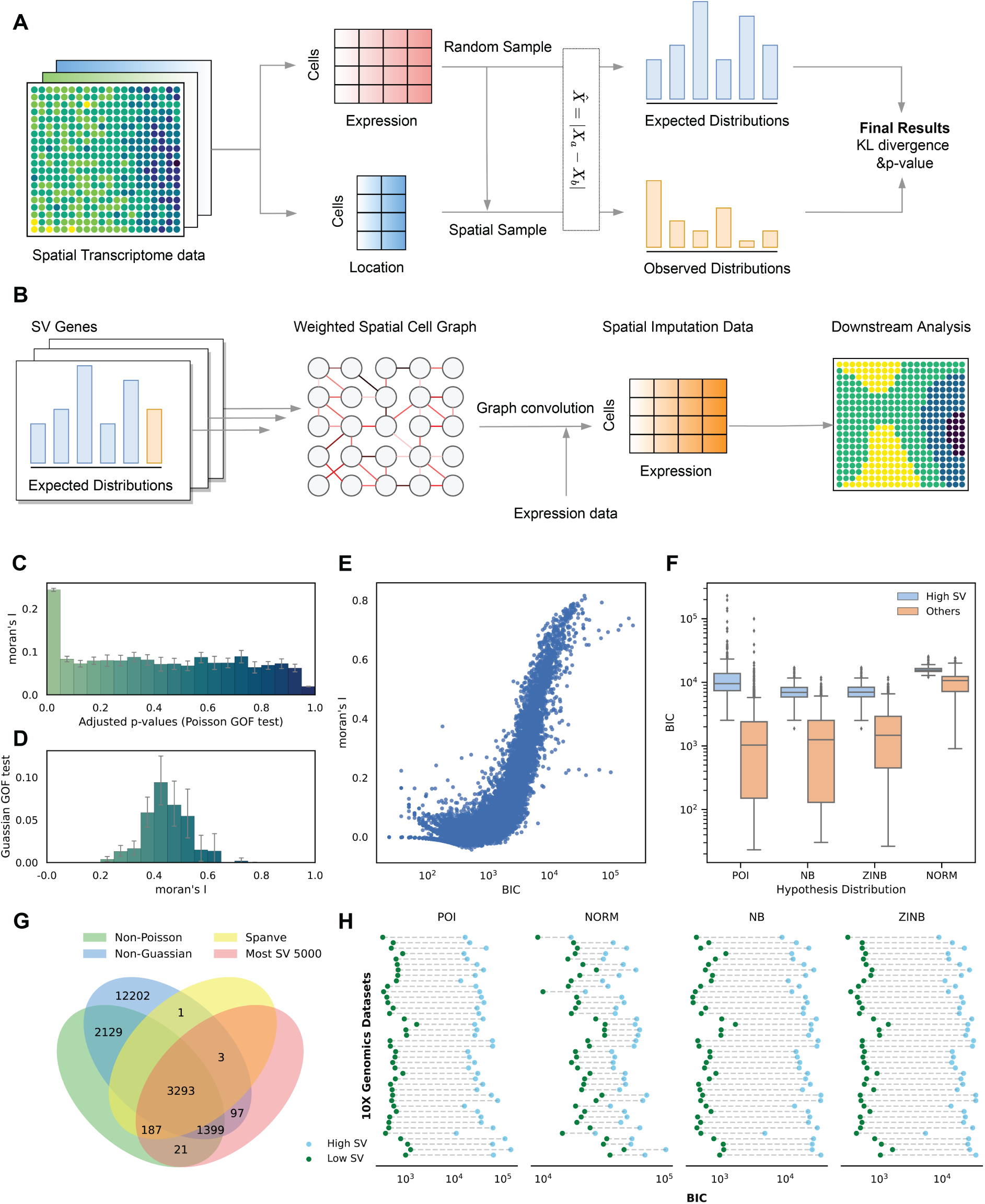
The framework and hypothesis of Spanve. (A) In spatial transcriptomic data, Spanve considers two types of sampling of cell or spot pairs: random sampling as the null hypothesis and location-based sampling as the alternative hypothesis. The spatial variability of genes is defined as the difference between two sampling distributions. (B) Spatial imputation employs graph convolution on a weighted spatial graph based on spot pairs that deviate from the expected distribution. Figure C to G depict one of the datasets from 10x Genomics. (C, D) The plots show the spatial variance, measured by Moran’s I, and the adjusted p values of the GOF test for the Poisson (C) and Gaussian (D) models. (E) The BIC value of the Poisson model shows a sigmoid-like relationship with the overall Moran’s I. (F) For four different distributions, genes with high Moran’s I values (> 0.5) tend to have higher BIC values. (G) The overlaps of non-Poisson, non-Gaussian, most SV 5000 genes and Spanve genes. (H) A general trend of high-SV genes with higher BIC values was observed in the 10x Genomics datasets.

These sampling approaches also aid in characterizing spatial domains by considering spot pair expressions during spatially variable gene identification. To smooth expression within spatial domains, we developed a spatial imputation method within the Spanve framework. This method identifies spot pairs with unexpected expression variations and constructs a network where edge weights represent the degree of deviation from expected expression differences between neighboring spots. The resulting network highlights unexpected differences in the overall dataset. A graph convolution strategy with self-loops (Figure 1B) helps mitigate technical errors and produces clustering-friendly imputed data.

To demonstrate Spanve’s advantages, we benchmarked its performance in spatially variable gene identification and downstream tasks, including spatial domain detection and spatial transcriptome deconvolution. We extended the classical spatial transcriptome analysis pipeline to include spatial pattern analysis and spatial co-localization analysis. Spatial pattern analysis groups spatially variable genes with similar spatial expression, helping researchers identify spatially-linked biological functions. Spatial co-localization generalizes spatially variable gene detection to gene pair detection, enabling exploration of cell-cell interactions and ligand-receptor relationships in a spatial context. These novel analytical approaches provide researchers with fresh insights into biological mechanisms.

Spanve is implemented in Python and can be easily utilized with minimal code. We provide a detailed protocol for Spanve usage within our extended analysis framework (Supplementary File 3).

## Results

### SV genes tend to violate the distribution assumption

As location effects exist, including cell type distribution, local environment and cell interactions, gene expression may not be described under simple distribution assumptions, whereas complex distributions largely increase the number of parameters and fitting time. Here, we showed that SV genes tend to violate the distribution assumption, revealing the inherent limitations of current statistical models based on assuming specific distributions. To show the relationship between the spatial variance represented by Moran’s I and the distribution assumption, the goodness-of-fit (GOF) test and Bayesian information criterion (BIC) were used to quantify how well gene expression can be described by the distribution. The anomalousness of gene expression is defined as the inability to model using a common hypothesis. We initially reported our observations for a single dataset (dataset identifier abbv. as VMOB, Table S2), subsequently extending our analysis to 43 additional datasets from 10x Genomics to show the generality of our conclusions (Figure 1C-H).

All genes were divided into 20 groups according to their Poisson divergence test *p* value. Genes with *p* values lower than 0.05 had a significantly greater Moran’s I than did the other genes (Figure 1C). After preprocessing, both the high-SV genes and low-SV genes differed from the Gaussian hypothesis (Figure 1D). We also compared the two test results with the spatial variance of the genes. Most SV genes could not be modeled considering the two hypotheses, while 3293 of 3297 genes identified by the Spanve model overlapped with the anomalous genes (Figure 1G). We then considered the NB and ZINB distributions. Overall, there was a sigmoidal relationship between the BIC score and Moran’s I (Figure 1E, Figure S1A). For high-SV genes, drop-out-like events caused by spatial effects may occur, and there is a small advantage over Poisson fitting (Figure 1F).

We then tested the generality of the findings using 43 datasets. The results showed that high-SV genes always had significantly greater BIC values when modeled by four distributions (Figure 1H). Most SV genes detected by various methods exhibited distribution anomalies (Figure S1B). Notably, Spanve achieved the highest proportion of anomalous SV genes while identifying the lowest number of SV genes (approximately 20% of all genes), which is consistent with a recent study ^15^ showing that most SV detection methods are overly sensitive, thus assigning a majority of genes as SV genes. This result highlights the specificity of the Spanve method, which outperforms other methods in identifying a more specific set of SV genes. In summary, our observations have shown that assuming specific distributions may result in a high false-positive rate for SV gene detection, and avoiding the distribution assumption may increase the specificity of the Spanve.

### Spanve decreases the number of false-positive outcomes with efficient large-scale computing

Then, we conducted comprehensive benchmarking (Figure 2A) of 11 SV gene identification methods, which can be roughly categorized into Gaussian process regression-based methods, clustering-based methods, spatial autocorrelation methods and others (Table S1). As there is currently no established gold standard for SV gene identification, we employed two global spatial autocorrelation metrics, Moran’s I and Geary’s C ^13^, to assess the statistical sensitivity of each method. Our benchmarking analysis was performed on 43 datasets (Table S2), which were preprocessed by Li et al. ^16^. For methods such as the sepal and silhouette coefficient rank in Giotto (Giotto-siRank), which only scores genes, the top 10% of genes were assigned as SV genes. The results (Figure 2C) showed that Spanve achieved the best score for both Moran’s I (0.41 ± 0.13) and Geary’s C (0.59 ± 0.13) while also identifying a moderate number of genes (95% confidence interval, 198 to 430). For the two different sampling strategies of Spanve, the K-nearest neighbor (KNN) network contains more edges than does the Delaunay network and is thus more specific to spatial variability but requires more computations. The improved performance in identifying SV genes is likely attributed to the reduced ratio of false positives, as the number of identified SV genes also decreased.

**Figure 2.**
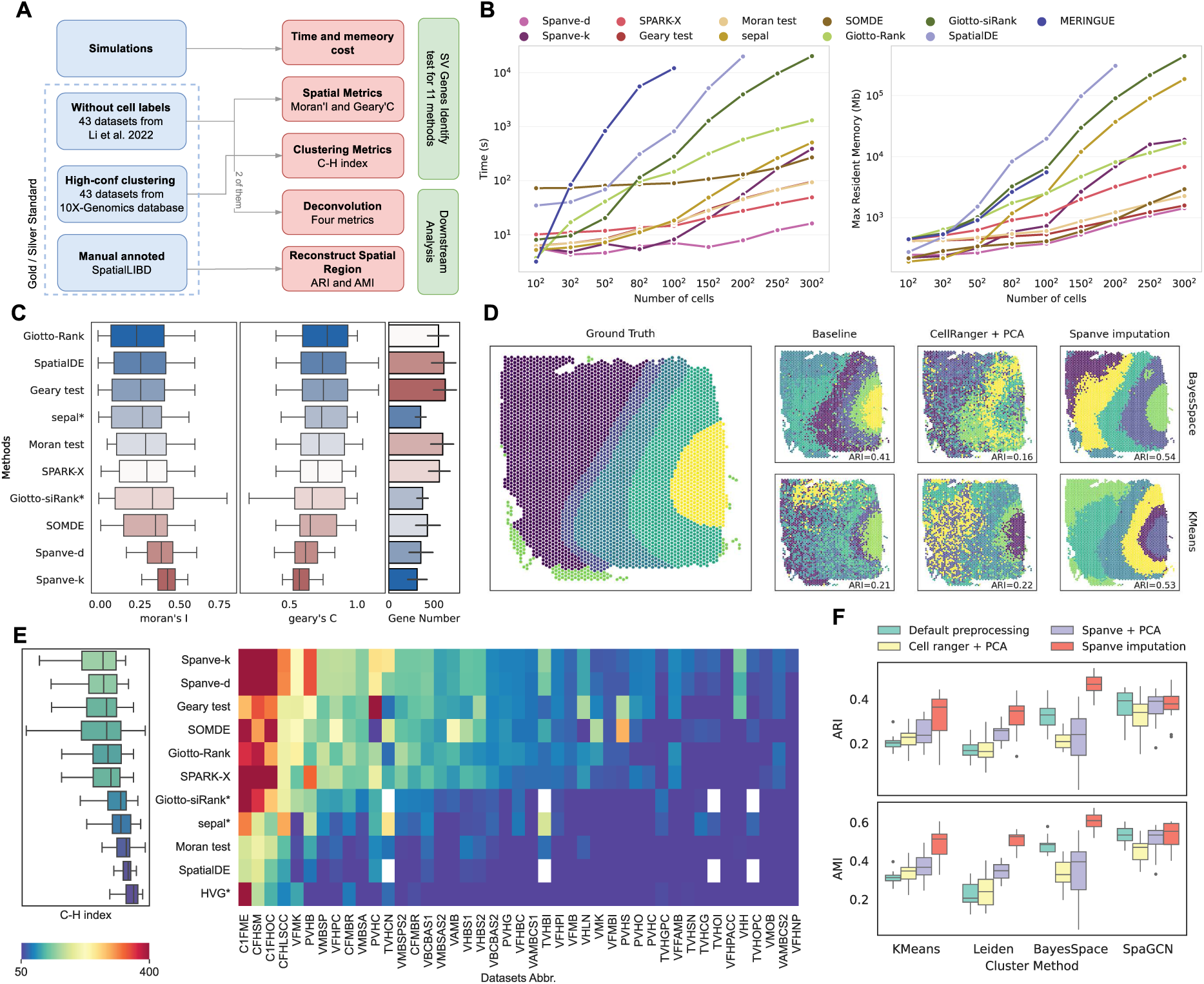
Evaluations of Spanve and 10 other methods. (A) datasets and evaluation across four aspects: computational cost, detection of spatially variable (SV) genes considering spatial heterogeneity and clustering awareness, and downstream tasks. (B) Time (left) and memory (right) costs for 1000 genes as cell numbers increase. (C) Spatial heterogeneity of detected SV genes. Box plot elements: centerline, median; box limits, upper and lower quartiles; whiskers, 1.5x interquartile range; color, performance. Bar plot error bars represent 95% confidence intervals. (D) Example of Spanve imputation aiding tissue structure identification (DLPFC dataset No. 151672). (E) Cluster awareness of SV genes in 43 datasets. Methods ranked by the median C-H index across datasets. Methods with stars only score genes without p values. Box and heatmap colors represent scores, and white indicates that no SV genes were detected. (F) Clustering results of the DLPFC dataset (n=12) with different preprocessing methods for the four spatial domain detection methods.

We then focused on estimating the computational cost of each method. The time and memory required to process the simulation data from a Poisson distribution with 1000 genes and varying numbers of cells were estimated. Each method was repeated thrice for each sample size to obtain stable results. The results (Figure 2B) showed that Spanve-d was the most efficient method, completing the analysis in 16 seconds and using 1500 megabytes of memory on data with up to 90000 cells. Spanve-k is fast and saves memory when the number of spots is less than 10000, but its performance deteriorates as the number of edges in the KNN network increases exponentially with the number of spots. These findings demonstrate that our method is scalable and suitable for the analysis of large-scale spatial transcriptome data.

### Utilizing the Spanve aids in delineating spatial domains

Spanve considers the complicated interactions between spots in the detection of spatial patterns, which is helpful for identifying the next crucial downstream task of spatial domain identification. To benchmark the relationship between clusters and SV genes, 43 datasets from the 10x Genomics database were utilized. These datasets include cluster labels, which were assigned using Space Ranger. The Calinski‒Harabasz (CH) index was used as an indicator to demonstrate how well selected genes are related to clustering results by the dispersion of data between clusters and within clusters. The higher the CH index is, the more likely it is that SV genes reflect clustering labels, implying that SV genes are more representative of the raw data. Our study involved a comparison of the CH indices of genes identified through various approaches, thereby highlighting the superiority of Spanve in recognizing cluster-aware genes (Figure 2E).

We also determined whether the Spanve imputation method could help with spatial domain detection or clustering. To validate whether imputation is beneficial for clustering, the spatial imputation process was aggregated with several clustering methods on a manually annotated human dorsolateral prefrontal cortex (DLPFC) ST dataset ^17^, which contains 12 samples. The four clustering technologies used include K-means, the Leiden algorithm, BayesSpace ^18^ and SpaGCN ^19^. BayesSpace is a clustering method that uses a Bayesian nonparametric model, and SpaGCN is a deep learning clustering method that integrates graph convolutional networks and ST data to learn the data cluster structure. We evaluated the performance of each clustering approach with different preprocessing methods and compared the resulting clusters with manual annotations by two clustering metrics, the adjusted Rand index (ARI) and adjusted mutual information (AMI), which evaluate the consistency of the predicted label and ground truth. For preprocessing, we considered the default processing of cluster methods, gene selection with principal component analysis (PCA), and Spanve imputation. For gene selection, Cell Ranger’s HVG selection and Spanve SV gene selection were included. We first chose one sample to visualize (Figure 2D) the K-means and BayesSpace clustering results, indicating that a clearer boundary of tissue layers can be obtained with Spanve imputation. Then, the performances on all 12 samples in the DLPFC dataset were compared (Figure 2F). With K-means, Leiden and BayesSpace, there were large improvements in the ARI and AMI with Spanve imputation. Notably, SV selection with PCA outperformed HVG selection, providing further evidence that SV genes help to characterize spatial domains. For SpaGCN, as we recognize that SpaGCN performs clustering on a graph, spillover imputation may make use of spot similarity for clustering and thus has little impact on it.

### SV genes facilitate spotwise cell type deconvolution

As each spot in an ST dataset may contain transcripts from multiple cell types, deconvolution algorithms are applied to estimate the proportional contribution of each cell type to the overall gene expression profile observed in a spot. In this section, we demonstrate that Spanve may also aid in cell composition inference. The rationale for leveraging SV genes for deconvolution lies in the assumption that the spatial distribution of these genes is correlated with underlying cellular heterogeneity. To accomplish this, we employed a previously reported standard benchmark ^16^ utilizing single-cell resolution ST data from the seqFish ^20^ and STARmap ^21^ datasets. These datasets allowed the generation of synthetic ST data with a known ground truth of the cell density at each spot. We evaluated the performance of three deconvolution tools (Cell2location ^22^, Tangram ^23^ and BayesTME ^24^) in conjunction with six gene selection methods and six combinations that integrate HVGs, marker genes of cell types, and SV genes identified using three different methods (Spanve, Moran’s I and SpatialDE). Overall, the performance was assessed using four metrics that were combined into a rank score. Details of the deconvolution estimation are included in the STAR Methods (Figure S1C–S1D).

The results demonstrated that proper gene selection improved the deconvolution performance compared to that using all genes (Figure 3A). Among the six gene selection methods, Spanve, along with its detected SV genes, achieved the best overall performance and was the only one that outperformed all the genes. For the combination of genes, incorporating SV genes can boost the performance of all three deconvolution methods. In particular, the incorporation of Spanve genes resulted in the greatest enhancement of deconvolution accuracy (Figure 3A-3B). This superior performance can be attributed to the ability of Spanve to detect the most relevant genes for cell type identification. We sought to determine whether genes that exhibited a high degree of correlation with specific cell types could be identified using SV gene detection methods. In the seqFish dataset for astrocytes, we found that Spanve identified 5 out of 10 such genes, SpatialDE identified 2 out of 10 genes, and Moran’s I identified 0 out of 10 genes (Figure 3C). By conducting an in-depth analysis of the distinct genes identified by the three spatial methods, it was evident that Spanve demonstrated the greatest overlap with the most crucial genes (Figure 3E) and exhibited the highest layer correlation (Figure 3C). Although most existing methods overlook the contribution of SV genes, our findings underscore the importance of incorporating these genes to facilitate accurate spotwise cell type deconvolution. The Spanve framework capitalizes on this opportunity by identifying cell type-relevant SV genes, thereby enhancing the deconvolution process and enabling a more comprehensive characterization of cellular compositions within the ST data.

**Figure 3.**
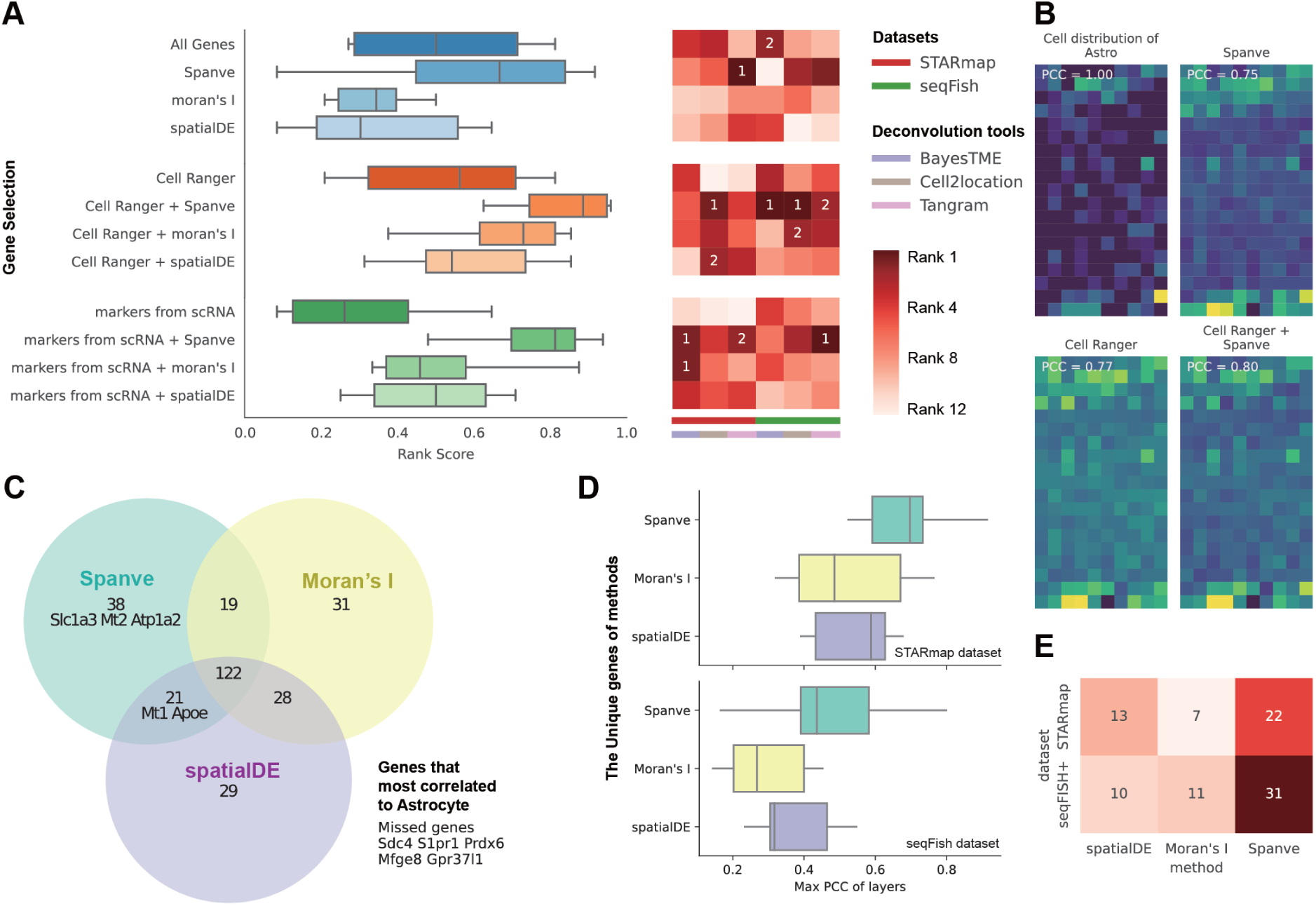
The Spanve enhances ST data deconvolution. (A) Different gene selection methods impact the deconvolution results across datasets and tools. The heatmap ranks the methods, labeling the top 2 performers in each run. (B) Astrocyte cell density and deconvolution results using different gene selection methods. (C) Venn diagram showing the overlap of SV genes detected by three methods in the seqFish dataset. The top 10 genes with the highest Pearson correlation coefficient (PCC) to astrocyte cell density are labeled. (D) Maximum PCC with ground-truth cell distributions for distinct genes identified by each of the three methods. (E) Number of distinct genes from each method overlapping with the most correlated genes, which include the top 30 genes for each cell type.

### Characterization of intratumoral heterogeneity by Spanve

A breast cancer dataset was used to demonstrate how Spanve helps in the analysis of ST data and identifies spatial microenvironment characteristics. We collected spatial transcriptomic data of human breast cancer tissue from the 10x Genomics Visium platform. This dataset provides whole transcriptome expression profiles and spatial coordinates of 4989 spots from a fresh-frozen tissue section derived from a patient with invasive ductal carcinoma, the most common type of breast cancer.

We designed a pipeline to illustrate how Spanve is employed in ST analysis (Figure 4A), including spatial domain detection, deconvolution, spatial pattern identification and spotwise cell‒cell interaction. At the beginning of our analysis, we applied Spanve for the identification of 4025 SV genes and imputation. Spanve imputation can aid in clustering by smoothing gene expression within domains while preserving the boundaries of spatial domains with differences, as shown in Figure 4B. Subsequently, the K-means algorithm clustered all the spots into seven spatial domains (Figure 4C), with the number of clusters determined by the elbow method based on the BIC. To determine the tissue type of the seven spatial domains, we utilized a single-cell resolved atlas of human breast cancers as a reference ^25^ to infer the cell composition of each spot and listed the top three main cell types in each domain in Figure 4D. Enrichment scores are the ratio of the number of cells in the spatial domain compared to that in the whole tissue, indicating whether a cell type is enriched (score > 1) or depleted (score < 1). We also compared the spatial domain detection results obtained using different methods based on deconvolution results and manual annotations. We found that other methods may fail to delineate the tumor region with the surrounding immune layer (Figure S2), whereas under the Spanve framework, K-means also obtained reasonable spatial domains with clear boundaries. To better understand the biological signals underlying SV genes, we performed a spatial correlation clustering method to group SV genes into several patterns (see STAR methods for details), followed by a commonly used gene set scoring method to show the consensus expression of spatial patterns (Figure 4E).

**Figure 4.**
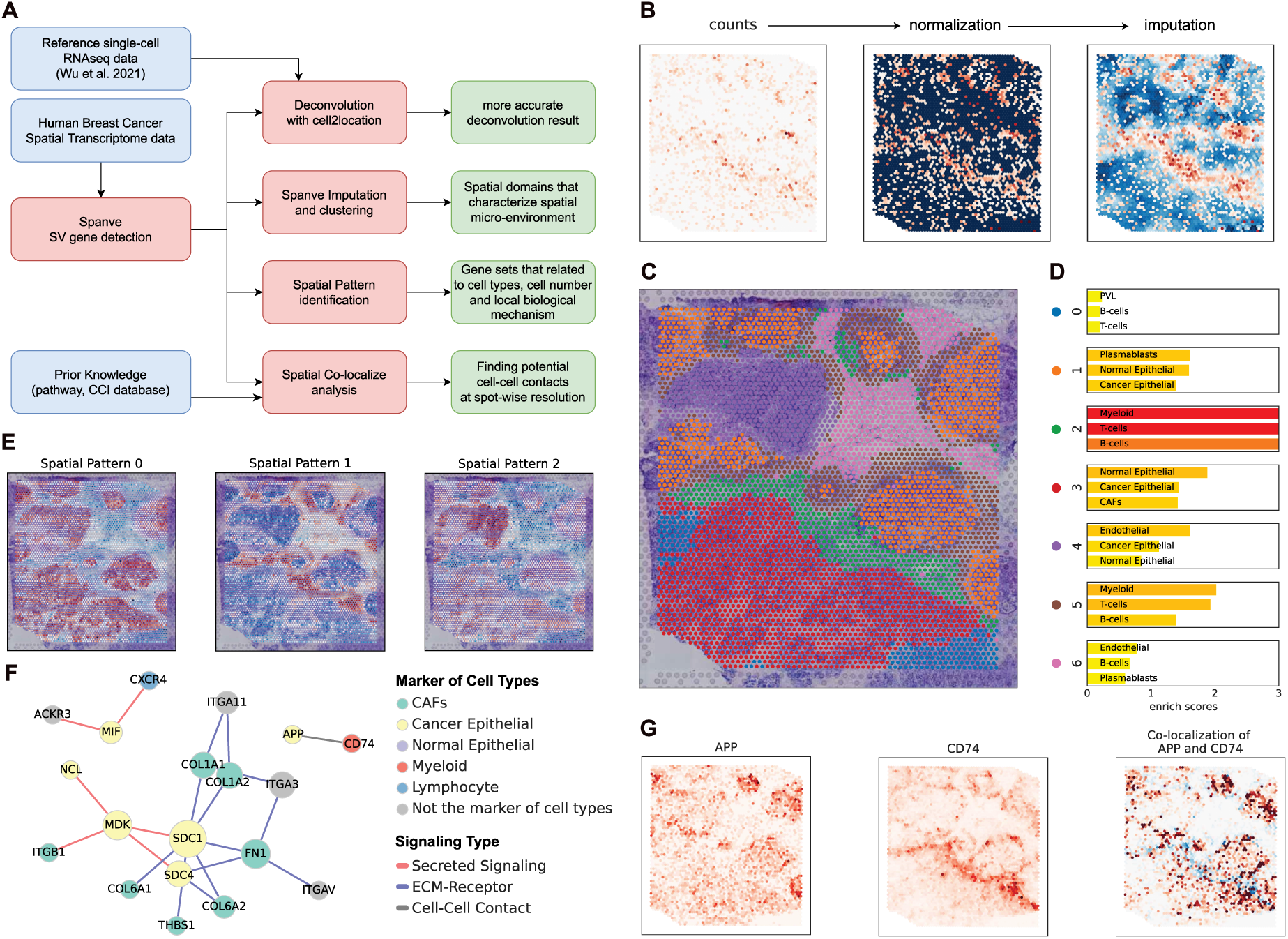
Analysis of human breast cancer ST data using Spanve. (A) Analysis pipeline under the Spanve framework. (B) Spanve imputation enhanced the spatial expression pattern of the CD74 gene. (C) Clustering result obtained using Spanve imputed data. (D) Main cell types identified in each cluster. The bar width and color represent the enrichment score, indicating over- or underrepresentation of the cell type in the cluster compared with the global distribution when the score is above or below 1, respectively. (E) Three distinct spatial patterns detected by Spanve. (F) Colocalization network of cell‒cell contact genes. Node colors represent cell types in which the genes are highly expressed. Edge colors denote signaling types associated with colocalized gene pairs. (G) Colocalization pattern of the genes APP and CD74.

Interestingly, visualizing the consensus expression of each pattern showed that spatial patterns were highly correlated with the spatial densities of some cell types. Thus, we linked the patterns to the factors and assigned a potential source of SV (Figure S3, Figure S4A).

Overall, these can be categorized into three types: those related to cell number, those caused by unbalanced cell type spatial density, and those caused by local biological mechanisms, which have small correlations with each other but focus within a spatial domain. To validate whether genes in the spatial pattern can show the spatial variance source, we checked the genes in spatial pattern 0 (which is related to the cell number) against the housekeeping genes from the Housekeeping and Reference Transcript (HRT) atlas ^26^ and found a large overlap (350 of 722, chi-square *p* value < 0.001). Thus, we could also perform gene set enrichment analysis to match biological functions to spatial patterns (Figure S4B).

We then investigated whether spatial colocalization could reveal the spatial characteristics of cancer tissue. We extended our algorithm to test whether a gene pair exhibits a spatial pattern, specifically by transforming coexpression into count-like data, inspired by the Pearson correlation formula. Spatial colocalization may indicate short-distance cell‒cell interactions (CCIs) at the resolution of a spot, which is often ignored by most current studies that focus on cell communication between spot clusters. We first checked the gene pairs from the CellChat DB ^27^ and displayed the significant gene pairs in a network (Figure 4F). We then detected a relationship between the CCI and CD74 and APP levels, which may exhibit a pattern correlated with the tumor region (Figure 4G). CD74 is a cell surface receptor that is involved in the formation and transport of MHC class II peptide complexes and is highly expressed in myeloid cells, B cells and endothelial cells according to reference scRNA-seq data. This cell interaction has been shown to be related to cancer development and immunotherapy in bladder cancer ^28^. Thus, the spatial variance in colocalization may indicate intratumor heterogeneity in the sensitivity to immunotherapy.

We also collected the seven most commonly studied pathways in breast cancer and performed a spatial colocalization test for each gene pair in the pathways (Figure S4C). The WNT signaling pathway and immune-related pathway were found to be the most significant gene pairs. Some of the most significant gene pairs are shown in Figure S4C. An interesting gene pair was TNFB3 with BAMBI, which showed tumor heterogeneity, especially in Cluster 4. We noticed that this gene pair plays a role in the TNF-beta signaling pathway, suggesting that it may be correlated with tumor malignancy. Compared to other tumor regions, a marker SV gene of Cluster 4 is LINC00645, which has been shown to induce epithelial– mesenchymal transition via TNF-beta in glioma ^29^. Based on the specific colocalization of the gene pair and the marker of Cluster 4, we assigned this strain to be invasive.

For validation, the top 15 marker genes of Cluster 4 were used to determine the relationship between survival and these genes. By selecting only ductal carcinomas in the TCGA-BRCA project, the expression of these genes was significantly related to the progression-free interval (PFI). Interestingly, during the first 500 days, there was no significant difference between the two groups, whereas the hazard probability dramatically increased after more than 500 days (Figure S4D). This result supports our assumption. These findings demonstrate that Spanve can be used to dissect spatial heterogeneity within a tumor sample, elucidate microenvironmental factors influencing cancer progression, and provide insights into potential cell‒cell interactions and signaling events.

## Discussion

In this study, we aimed to find a potential solution for balancing false-positive rates and computational costs. Our findings showed that SV genes tend to be difficult to model using normal, Poisson, NB, and ZINB distributions. As a result, current methods that depend on an expression distribution hypothesis may yield false-positive outcomes if the genes do not follow this hypothesis. To address the abovementioned challenge, a spatially variable gene identification method named Spanve has been proposed.

Our method provides very close insight into Geary’s C, which also measures the differences in expression between neighboring spots. Notably, our method reduces the false-positive ratio by considering the deviation of the overall distribution, whereas Geary’s C only uses it to normalize the metric. In this respect, it can be thought that Spanve takes into account the background distribution. Some studies have attempted to modify Geary’s C, but false-positive outcomes were not considered ^30^. Due to the improved specificity, Spanve boosts downstream tasks in ST analysis by better feature selection ability and a spatial imputation method, which can be integrated into existing analysis pipelines. A case study on a breast cancer sample showed that it is possible to characterize the microenvironment of the tissue under the Spanve framework. Spatial coexpression has been considered by some previous studies ^31–34^, incorporating cluster or spatial domain information in the detection process. In contrast, our study proposes a spotwise gene colocation or coexpression method, enabling the quantification of coexpression at the spatial location level. This approach may be important when considering local biological mechanisms that exhibit specificity within discrete spatial domains.

In addition to these benefits, the scalability of Spanve will become increasingly crucial as the volume of ST data expands. Compared with current methods, the Spanve method requires less time and has a lower computational cost for large-scale spatial transcriptomic data. The computational complexity of the Spanve algorithm depends on the data complexity, with sparser data leading to simpler counting probabilities. As the volume of spatial spots increases, they are expected to become increasingly sparse, as shown by single-cell sequencing ^35^. Therefore, Spanve may be of increasing importance when higher-resolution ST technology appears.

### Limitation of the study

The current Spanve approach presents several limitations that warrant consideration. One key limitation is its inability to simultaneously process data from similar tissue slices to detect more robust SV genes and leverage prior knowledge. The integration of prior data or knowledge could potentially help identify more robust SV genes and uncover strongly related or even causal biomarkers, which would be particularly valuable for single-dataset studies. In future work, this issue might be addressed by modifying the spatial sampling strategy to incorporate prior data and knowledge.

Another limitation is that Spanve is designed specifically for transcriptomics data, as its sampling method is most suitable for discrete data. This specificity may restrict its application in the rapidly evolving fields of spatial proteomics and spatial multi-omics. To extend the nonparametric statistical framework to discover spatial variance in other modalities, it may be necessary to develop appropriate methods for converting continuous data into discrete formats.

Furthermore, the analysis of spatial patterns and spatial colocalization presents validation challenges. Some identified spatial patterns may not readily correlate with specific biological functions, local biological processes, or cell type differences, making interpretation challenging. Additionally, the current analysis framework may not be suitable for detecting long-distance cell-cell interactions, such as those mediated by secreted factors. This limitation restricts the applicability of the spatial colocalization analysis to short-range interactions. Addressing these limitations in future iterations of Spanve will enhance its utility and broaden its applicability across various spatial omics technologies and biological questions.

Overall, the Spanve method represents a significant advancement in ST analysis, providing researchers with a powerful tool to extract meaningful insights from spatial gene expression within biological samples. The improved specificity, enhanced downstream utility, and computational scalability of Spanve position it as a valuable addition to the ST analytical toolkit.

## STAR Methods

### Key resource table

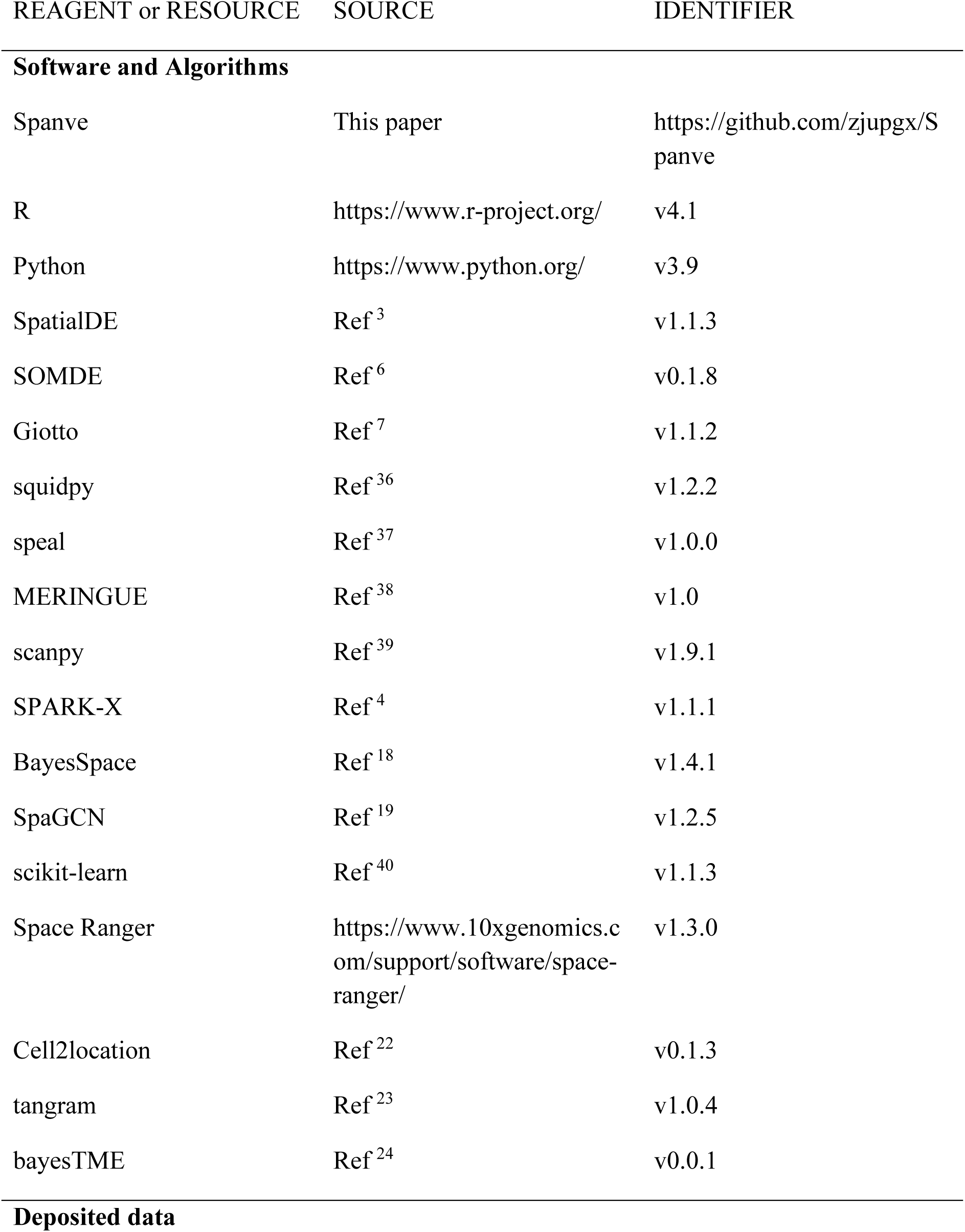
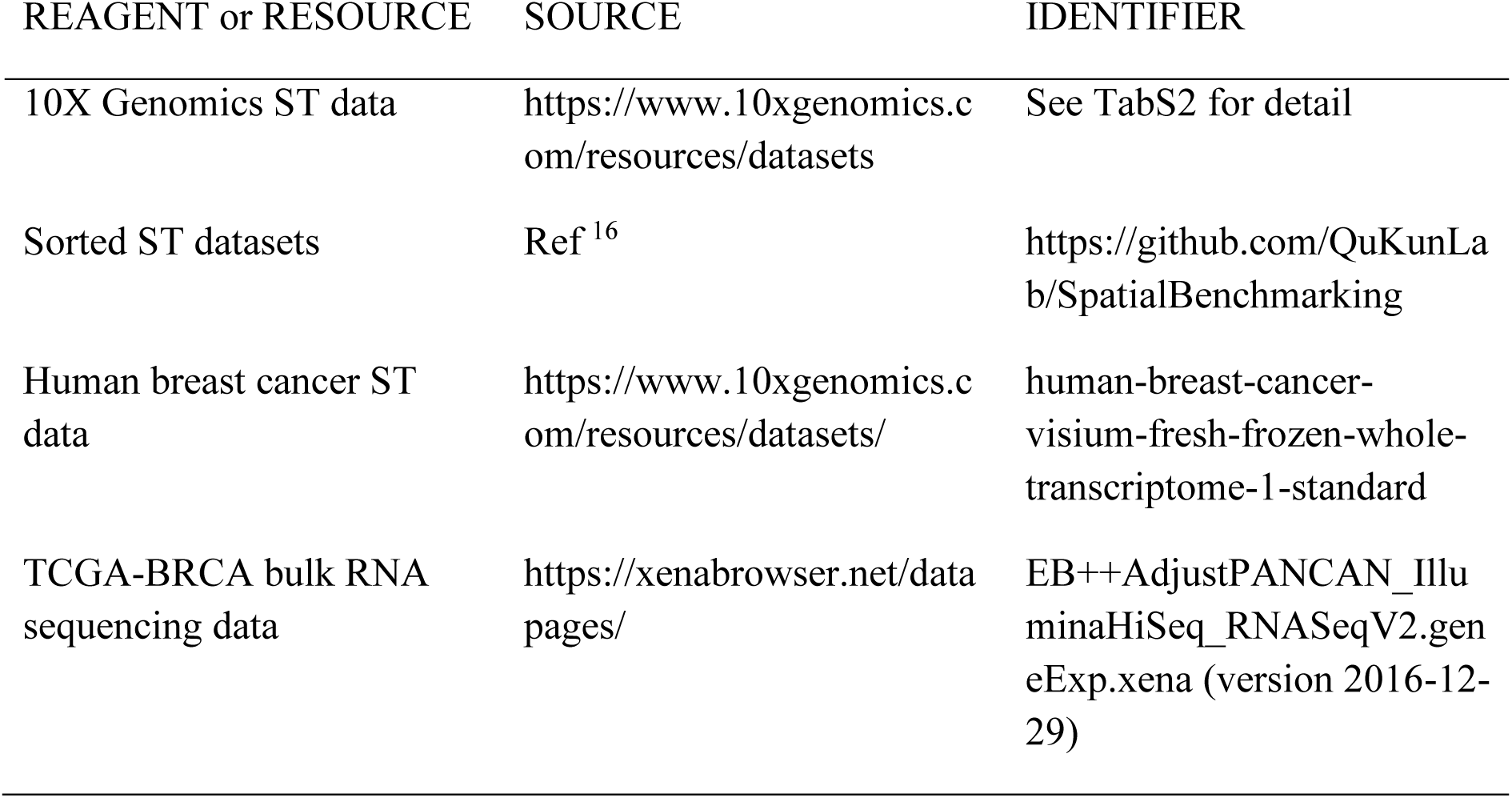

### Methods details

For the spatial transcriptome raw count data *X* ∈ ℕ^*N*×*M*^ with *N* sequencing spots and *M* genes and any gene expression *x* ∈ ℕ^*N*^, the two-dimensional location *L* ∈ ℝ^*N*×2^ of each spot is also obtained. The aim of SV gene identification is to identify genes whose expression is dependent on spatial locations *x* ⊥ *L*. From an intuition of spatial variance that can be treated as the difference between cells and their neighbors, Spanve uses absolute subtraction to evaluate the difference and transforms the dependence problem into a contrast of two samplings, one null sampling, and the other spatial sampling. Three steps are involved in the identification of SV genes: 1) from observation to build a null sampling distribution, 2) build a spatial network and perform spatial sampling, and 3) take a statistical test for the spatial variance.

#### ST data preprocessing

Spanve takes spatial transcriptomic data as inputs, which consists of two parts: the raw count expression of genes and locations. Although preprocessing may play an important role in denoising spatial transcriptomes, a special strategy is used, where the median of the data remains consistent for each gene:

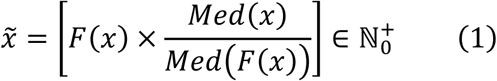

where *x̃* is the preprocessed data, *F* is any of the preprocessing functions, [.] transforms the variable into a discrete value and *Med* calculates the median of the data. It is notable that preprocessing is not a necessary step for Spanve, and a worse result may be obtained if data have only parts of the genes or if some cells have very low counts. In this study, preprocessing was performed only when the data were undivided.

#### Sampling Strategy

We aim to quantify the spatial effects of gene expression by measuring differences in the expression levels of the two cells. A simple and intuitive way to do this is to use the absolute subtraction difference (ASD) of any two cells, which reflects the magnitude of their expression discrepancy, regardless of the direction. To calculate the ASD of any two spots, we define a new matrix *X̂* ∈ ℕ^*N*×*N*×*M*^, where each entry *X̂_ij_* is a vector of length *M* that contains the ASD values for each gene between cell *i* and cell *j*. Formally, we have

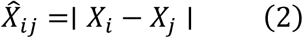

where *X*_*i*_ and *X*_*i*_ are the rows of *X* corresponding to cell *i* and cell *j*, respectively. Note that obtaining the entire matrix *X̂* is not scalable, and our sampling strategy only considers a part of the matrix. In this step, our goal is to compare the distribution of ASD values for each gene under two different scenarios: spatial and random sampling. Spatial sampling means that we only consider cell pairs that are close to each other in physical space, whereas random sampling means that we consider all possible cell pairs regardless of their spatial proximity. By comparing these two distributions, we could assess whether the expression of a gene is influenced by its spatial location. For each gene *g*, we denote the distribution of ASD values under spatial sampling as *P_obs_*(*x̂_g_* | *x: L*) and under random sampling as *P*_exp_(*x̂_g_*), where *x̂_g_* is the vector of ASD values for gene *g* across all cell pairs. As the expression of all cells is observed, it is easy to obtain the expected random sampling result of the distribution of ASD *P*_exp_(*x̂_g_*) by listing all combinations.

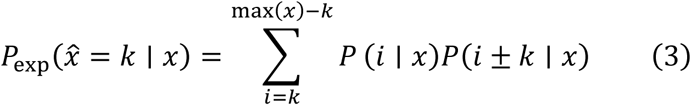

The spatial sampling in this study is based on a spot network that represents the spatial proximity of spots *S* = {*S*_*n*_, *S*_*e*_}, where nodes *S*_*n*_ are cells and edges *S*_*e*_ indicates that the two cells are sufficiently close. The edges in the network represent sampled spot pairs. We use two types of networks: K-neighbor and Delaunay networks. The K-neighbor network connects each cell with its K-nearest neighbors based on Euclidean distance (by default, the number of neighbors is set to [*N*/100]), whereas the Delaunay network connects each cell with its neighbors to form a Delaunay triangulation. A Delaunay network is a type of triangulation used to model data points where no point is inside the circumcircle of any triangle. The Delaunay network has shown scalability as the number of edges in Delaunay networks (< 3*N* − 3) is smaller than that in KNN (*N* × [*N*/100]). Based on our research, we have determined that the optimal method for constructing a network is contingent upon the number of available spots. The KNN network of spatial location is used when the sample size *N* < 10000; otherwise, the Delaunay network is used. By calculating the ASD values for all cell pairs connected by an edge in the network, we obtain the distribution of the spatial sampling *P_obs_*(*x̂_g_* | *x: L*) for each gene.

#### Statistics of spatial variance

Here, we take the Kullback-Leibler divergence, *D*_*K*_*_L_*, to quantify the difference in the distribution:

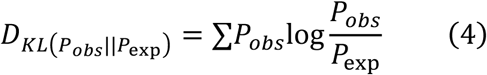

To determine how large a KL divergence can be considered to be spatially variable, we determined the threshold by a modified G-test process. As the expression counts of genes are discrete, the ASD value is also discrete. We treated *x̂* as a categorical variable, and its possible value, that is, the degree of freedom, is from zero to the maximum expression max(*x*). The KL divergence can be related to the G test, where G statistics are proportional to *D*_*KL*_ ^41^.

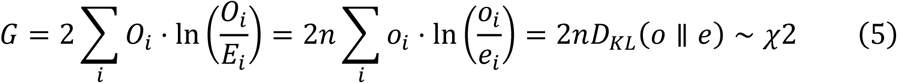

where *n* is the total number of observations *n* = ∑*O*_*i*_. However, having strong weights between neighbors also means that those locations provide “redundant” information, since neighbors tend to be similar, and they do not provide independent data points. If the expression of one of the spots changes, each edge link to the spot will change, that is, The ASD value here is not independent. Consequently, we modified the formula of the G-statistic to avoid false positives by replacing *n* (in our methods, it represents the number of all pairs) with the number of spots *N*. Intuitively, the effective sample size over all observations *x̂* i s *N* under the null hypothesis.

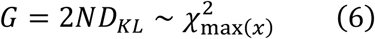

#### Spanve Imputation

The imputation of the Spanve is performed based on the previous SV gene results. Overall, it is a simple graph convolution method for a modified spatial network. We first modified the previous networks that reflect the spatial locations by adding edge weights as the fraction of SV genes that significantly (by default, with a confidence level *⍺* of 0.05) break the expectation. The new network can be viewed as a boost in spatial effects.

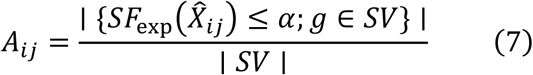

Then, graph convolution with a self-loop is performed under the expression matrix to obtain the imputation matrix *X*′ = *X*(*A*+ *I*)^2^, where *I* is the identity matrix.

#### Spatial patterns detection

The space matrix *S*_*A*_∈ ℝ^*N*×*N*^ is the affinity matrix of neighborhood network *S*. For genes *⍺* and *β*, *x*_*⍺*_ and *x*_*β*_ are the expressions. The space covariance is defined as 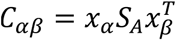, and the spatial correlation is 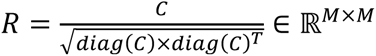. To construct a gene network, a correlation threshold is selected based on finding the maximum change in frequency, which is a heuristic approach. By Louvain algorithm, the gene network is divided into several gene community, or gene patterns.

#### Spatial co-localization

For any two gene *⍺* and *β*, let

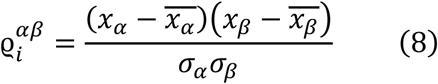

where *σ* is the standard variance of a gene and *x̄* is the mean of the expression. The mean of *C* was equal to the Pearson’s correlation coefficient *ρ*.

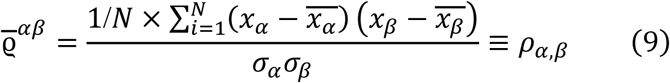

Thus, ϱ can be considered as the sample-resolution co-expression strength of the two genes. Thus, a similar strategy can be used to explore gene pairs with spatially variable co-localization. In this study, the pathway structure for the spatial co-localization test was collected from Pathway Commons ^42^. Prior knowledge of cell-cell interactions is collected from the CellChat DB ^27^.

### Computational cost evaluation

We simulated spatial transcriptomic datasets by assuming that gene expression follows a Poisson distribution, *PoiP*(*λ* = 5). A Gamma distribution (*k* = 2, *θ* = 5) is used to create gene-specific variability by scaling gene maximums to improve realism. All spots are regularly distributed in a square pseudo-splice, where the distances between the horizontal and vertical spots are equal (10, 30, 50, 80, 100, 150, 200, 250, and 300). Under each condition, all methods were run three times using different random seeds.

The computational cost was evaluated under the simulation data using the GNU time tool on a Linux server with Intel(R) Xeon(R) CPU E7-4850 v3 @ 2.20GHz (56 kernels) and 976 GB memory. All kernels are used if multi-thread is available.

### Benchmarking of spatially variable genes

Eleven SV identification methods were used here, which can be roughly categorized into spatial autocorrelation-based (Moran test, Geary test, MERINGUE), Gaussian process regression-based (SpatialDE ^3^, SPARK-X ^4^), cluster-based (SOMDE ^6^, Giotto-rank and Giotto-silhouette-rank ^7^), and others (sepal ^37^, Spanve-k, and Spanve-d). For nine of them, the adjusted p-value is the output, while for Giotto-silhouette-rank and speal, only the score is given. Thus, for score-only methods, the top 10% of score genes were treated as SV genes; for others, the threshold of the adjusted p-value was set to 0.05. For MERINGUE, the computational cost is too high to benchmark. The benchmarking runs on 44 datasets with only HVGs sorted by Li et al. ^16^. Furthermore, gene relationships with clustering results were scanned in silver standard datasets, which are the result of the Louvain algorithm in the decomposition of whole data. The CH index is used as an indicator to show how easy it is for genes to obtain a cluster label. With a higher CH index, the genes used as features were more closely related to cell types.

### Identification of spatial domains

Four clustering methods are used to identify the layer structure of the DLPFC: K-means (distance-based), Louvain (graph-based), BayesSpace (Bayesian-based), and SpaGCN (deep learning-based). Default parameters were used for benchmarking, and principal component analysis (PCA) embeddings used by the four methods were replaced with the PCA results of Spanve imputation. As K-Means clustering requires the parameter of cluster number, the elbow method based on inertia determines it. The inertia of K-means is defined as the sum of the squared distances of the samples to their closest cluster centers. For BayesSpace, there is no such method; thus, the ground-truth number of clusters is used. For SpaGCN, the tissue images were not included in the analysis. The gold standard for the tissue layers was manual annotation performed by Maynard et al. ^17^.

### Benchmarking of cell type deconvolution

For benchmarking cell type deconvolution, we followed a previous study ^16^ to perform and evaluate the performance. Specifically, two single-cell-resolution ST datasets were collected to generate multi-cell-resolution ST data. Based on spatial location, the data were split into grid-like sequencing spots containing multiple cells. By doing so, the simulated ST data and ground truth cell densities were obtained. Here, we use three deconvolution utilities, Cell2location, Tangram and bayesTME, to perform deconvolution based on the 12 different gene selection methods. For each gene selection method, 1000 out of 9684 genes from STARmap Dataset ^21^ and 200 out of 882 genes from seqFish Dataset ^20^ were selected based on the simulation. Spanve, Moran’s I, spatialDE, and Cell Ranger methods were applied to the ST data. Marker genes for single cells were selected using Student’s t-test on log-transformed counts, implemented by scanpy’s rank_genes_group. The most significantly differentially expressed genes were selected based on the sum of the t-statistics for each cell type. The combination of the two methods selects the union of genes that are detected by the two methods. Four metrics were used for evaluation: the Pearson Correlation Coefficient (PCC), Structural Similarity Index (SSIM), Root Mean Square Error (RMSE), and Jensen– Shannon Divergence (JSD). For each run, that is, a dataset with a deconvolution method and all gene selection methods, a rank score was calculated to combine the four metrics and highlight the differences. The rank score (AS) was defined as 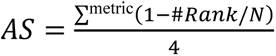, representing the average rank of each method across the four metrics.

### Analysis of breast cancer spatial transcriptomics

To perform a case study, we used public human breast cancer ST data available from 10x Genomics (Table S2). We initially performed quality control by removing cells with total counts less than 10 and genes expressed in fewer than 5 cells, obtaining 4898 spots and 20227 genes. The Spanve analysis framework was then used to obtain the results (Figure S4A). To determine the function of spatial patterns, GSEA was performed using library annotation from enrichr ^43^ by pygsea ^44^. To check the relationship between the top 15 markers of Cluster 4, we investigated the survival time of ductal carcinoma in preprocessed TCGA-BRCA data collected from xenahubs (https://xenabrowser.net/datapages/). The log-rank test was performed in the two groups by selecting the top or bottom 20% expression.

## Data and Code Availability

The source code of Spanve and reproducibility of the analysis can be accessed by FishShare ^45^ or Github (https://github.com/zjupgx/Spanve). The data underlying this article were all derived from sources in the public domain (See TabS2 for detail). 10X Genomics ST data can be achieved from https://www.10xgenomics.com/resources/datasets. Sorted ST datasets ^16^ are provide by authors: https://github.com/QuKunLab/SpatialBenchmarking. Human breast cancer ST data is downloaded from https://www.10xgenomics.com/resources/datasets/human-breast-cancer-visium-fresh-frozen-whole-transcriptome-1-standard. TCGA-BRCA bulk RNA sequencing pre-processed data is from https://xenabrowser.net/datapages/EB++AdjustPANCAN_IlluminaHiSeq_RNASeqV2.geneExp.xena (version 2016-12-29).

## Author contributions

G.X.C., X.G., and Z.Z. designed the study; G.X.C. implemented the method; G.X.C., Y.C.C. and S.Q.C. analyzed the data; G.X.C. and Z.Z. wrote the paper. Z.Z. supervised the work.

## Acknowledgements

We thank Drs. Binbin Zhou and Jingqi Zhou for their critical reading of the manuscript. We also appreciate the Information Technology Center and State Key Lab of CAD&CG, the Innovation Institute for Artificial Intelligence in Medicine, Zhejiang University for the support of computing resources.

