## Supplementary file 3 for "Spanve: A Statistical Method for Detecting Downstream-Friendly Spatially Variable Genes in Large-Scale Spatial Transcriptomic Data"

### Tutorial for exploring, explaining and analyzing SV genes under Spanve framework

Spatial transcriptomics (ST) is a powerful technique that allows researchers to study in situ gene expression, providing valuable insights into cell function mechanisms. One crucial aspect of analyzing ST data is the identification of spatially variable (SV) genes, which exhibit varying expression patterns across different spatial locations. Traditional methods for identifying SV genes often rely on modeling gene expression using distribution hypotheses, but this approach can lead to false positives. To address this limitation, a novel nonparametric statistical approach called Spanve has been developed. Spanve models spatial dependence as the distance between two sampling distributions without making any assumptions, resulting in lower false positives and facilitating more accurate downstream ST analysis.

In this tutorial, we will explore the Spanve framework, which provides a comprehensive solution for identifying, explaining, and analyzing SV genes in ST data. We will cover the theoretical foundation of Spanve, its implementation details, and its applications in various downstream analyses, including spatial domain detection, cell type deconvolution, and spot-wise cell-cell interaction analysis. By the end of this tutorial, you will have a solid understanding of how to leverage Spanve for your ST data analysis needs.

- How to install

```
pip install Spanve
```

And first, let's import the necessary libraries and load the dataset.

In [1]:
```
import os
import sys

import matplotlib.pyplot as plt
import numpy as np
import pandas as pd
import scanpy as sc
import seaborn as sns
import sklearn
import Spanve
import warnings

print(Spanve.__version__)
from Spanve import Spanve, adata_preprocess, adata_preprocess_int  # noqa: E402

warnings.filterwarnings("ignore")
sc.set_figure_params(dpi=150)
np.random.seed(2233)
```

```
0.1.4
```

#### Reading data and preprocess

In this section, we will load a spatial transcriptomics dataset from 10X Genomics called "Visium_Human_Breast_Cancer" and perform necessary preprocessing steps to prepare the data for analysis with Spanve.

First, we import the dataset using the `sc.datasets.visium_sge()` function from the Scanpy library, or you can download the dataset by the link. We then apply filtering steps to remove cells with fewer than 10 counts and genes expressed in fewer than 5 cell.

To prepare the data for downstream analysis, we create three new layers in the adata object:

1. "normalized": This layer contains the normalized gene expression values obtained by running adata_preprocess(adata).X.
2. "counts": This layer is a copy of the original count matrix, adata.X.copy().
3. "normlized_counts": This layer contains the normalized count values obtained by running adata_preprocess_int(adata).X.

Now, the adata object is ready for further analysis with Spanve. In the next section, we will explore how to identify spatially variable genes using the Spanve framework.

In [2]:
```
adata = sc.datasets.visium_sge('Visium_Human_Breast_Cancer')
adata.var_names_make_unique()
sc.pp.filter_cells(adata, min_counts=10)
sc.pp.filter_genes(adata, min_cells=5)
print(adata.shape)

adata.X = adata.X.toarray()
adata.layers["normalized"] = adata_preprocess(adata).X
adata.layers['counts'] = adata.X.copy()
adata.layers['normlized_counts'] =  adata_preprocess_int(adata).X
```

```
(4898, 20227)
```

#### SV genes detection by Spanve

After preparing the data, we can now run Spanve to identify spatially variable (SV) genes. Spanve requires count data as input, so we first need to assign the count matrix to adata.X. Alternatively, you can use the `adata_preprocess_int` function to obtain a preprocessed count matrix. But this process is not promised to perform a better performance.

Here, we've set n_jobs=16 to use 16 parallel jobs for faster computation. You can adjust this value based on your system's available resources.

Spanve has several important parameters that you can customize:

- K (int, optional): Number of K neighbors. Defaults to None, which means using adata.shape[0]//100 or 5.
- neighbor_finder (str, optional): Spatial graph construction method, one of ['knn', 'Delaunay']. Defaults to None. if number of samples < 10000, use knn else use Delaunay. For the difference of the two methods, please refer to the manuscript. Overall, the Delaunay method reduce the computational cost but knn get a more accurate result.
- n_jobs (int, optional): Parallel jobs. Defaults to -1 to use all available kernels.

The `svmodel.rejects` attribute contains a boolean array indicating whether each gene is identified as an SV gene or not. Summing this array gives us the total number of detected SV genes. Or you can check the `svmodel.result_df` for the detailed information of the SV genes.

In [3]:
```
adata.X = adata.layers['counts']
svmodel = Spanve(adata, n_jobs = 16)
```

```
svmodel.fit(verbose=True)
print("Detected SV gene number:",svmodel.rejects.sum())
svmodel.result_df.head()
```

```
#1 Expected Dist within Nodist Hypoth: 100%|██████████| 20227/20227 [00:05<00:00, 4007.77it/s]
#2 Nearest Neighbors Found
#3 Computing Absolute Substract Value: 100%|██████████| 20227/20227 [00:18<00:00, 1068.44it/s]
#4 Entropy Calculated
#5 G-test Performed
Write results to adata.var['spanve_spatial_features']
#--- Done, using time 25.91 sec ---#
Detected SV gene number: 4019
```

Out[3]:

| | ent | pvals | rejects | fdrs | max_expr |
|---|---|---|---|---|---|
| **AL627309.1** | 4.093391e-06 | 0.980150 | False | 1.0 | 2 |
| **AP006222.2** | 1.085609e-07 | 0.973985 | False | 1.0 | 1 |
| **LINC01409** | 9.909635e-07 | 0.995158 | False | 1.0 | 2 |
| **LINC01128** | 1.342611e-05 | 0.987803 | False | 1.0 | 3 |
| **LINC00115** | 1.757456e-06 | 0.991429 | False | 1.0 | 2 |

Saving the result to a file is also supported by the `save` method. By default, the result will be saved to a data frame.

In [4]:
```
svmodel.save('./spanve_model_result.csv')
# load the model
# loaded_model = svmodel.load('./spanve_model_result.csv')
```

#### Imputing ST Data with Spanve

Spanve can also be used to impute values in the spatial transcriptomics data, which can improve downstream analysis and interpretation. In this section, we'll learn how to perform imputation using the impute_from_graph method provided by Spanve. The normalized gene expression matrix is used as input. And the key parameters are:

- n_circle, int, optional: The number of circles to be used for imputation. Defaults to 2. Please note that the larger the number of circles, the more homologous the imputation expression. Not recommended to use a number larger than 2.

In [5]:
```
X = adata.layers["normalized"]
newX = svmodel.impute_from_graph(X, verbose=True, n_circle = 2)
adata.layers['imputated'] = newX
```

```
Impute data, there are 2 circles
generate graph from spatial genes: 100%|██████████| 4019/4019 [00:10<00:00, 372.55it/s]
Impute data, there are 1 circles
```

Here, we also show a gene impuation example.

In [6]:
```
f, axes = plt.subplots(1,3, figsize = (12,4))
for layer, ax in zip(['counts','normalized','imputated'], axes):
    sc.pl.spatial(
        adata, color = ['CXCR4'],
```

```
        vcenter=0, colorbar_loc=None,
        cmap = 'RdBu_r', layer=layer,
        img_key=None, spot_size=300,
        ax = ax, title = layer, show=False,
    )
    ax.set_xlabel('')
    ax.set_ylabel('')
    ax.axis('off')
```

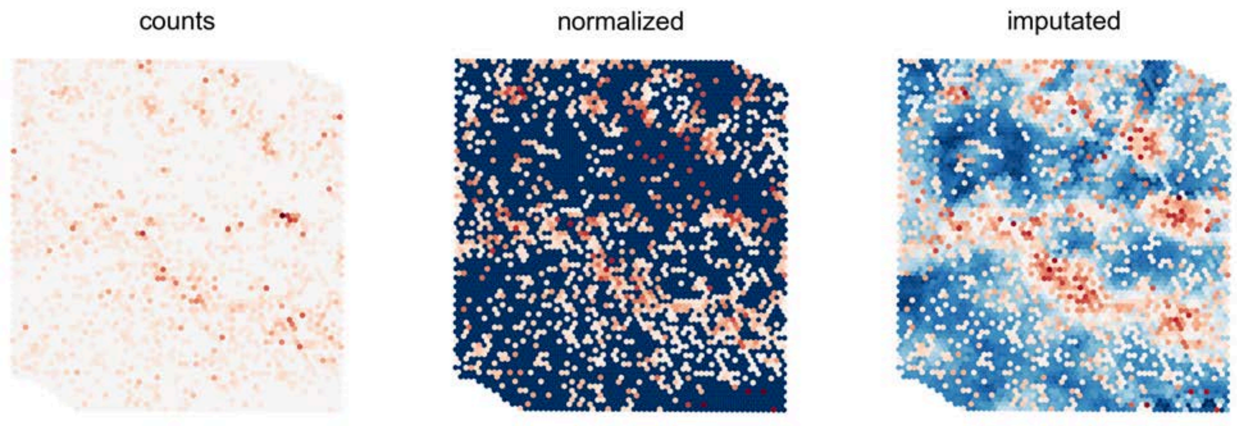


#### Exploring SV Patterns

After identifying the spatially variable (SV) genes, Spanve provides a way to explore their consensus expression patterns, known as SV patterns. These patterns are obtained by clustering the SV genes based on their spatial expression correlations, revealing groups of genes with similar spatial distributions. Spanve offers two different clustering methods for this purpose: Louvain clustering and Spectrum clustering.The detect_spatial_pattern method accepts several arguments to customize the clustering process:

1. clustering_method (str, optional): 'networkx' (Louvain clustering) or 'sklearn' (Spectrum clustering). Defaults to 'networkx'.
2. n_clusters (list, None, or int, optional): Only used when clustering_method is 'sklearn'. Defaults to None. If None, it will search for the optimal number of clusters between 3 and 10. If a list is provided (e.g., [3, 10]), it will use AutoCluster to search for the best cluster number within the specified range.
3. search_genes (list, optional): A list of gene names to consider for SV pattern detection. Defaults to None, which means all SV genes will be used.
4. aff_thres (float, optional): The threshold for the edge weight in the correlation graph. Defaults to None, in which case the threshold will be set to the value where the frequency of spatial correlation is highest.

After running svmodel.detect_spatial_pattern, Spanve will detect the SV patterns and store the results in the svmodel.result_df attribute. You can then use various visualization and analysis methods provided by Spanve to explore and interpret these patterns.

In [7]:
```
pats = svmodel.detect_spatial_pattern(
    verbose=True,
    clustering_method = 'networkx', # or 'sklearn'
    resolution = 1.25 # kwargs pass for the clustering method
)
```

```
using networkx algorism get communities : 10
 With size:  [ 200 1603   30  132   69  326  109 1392  148   10]
```

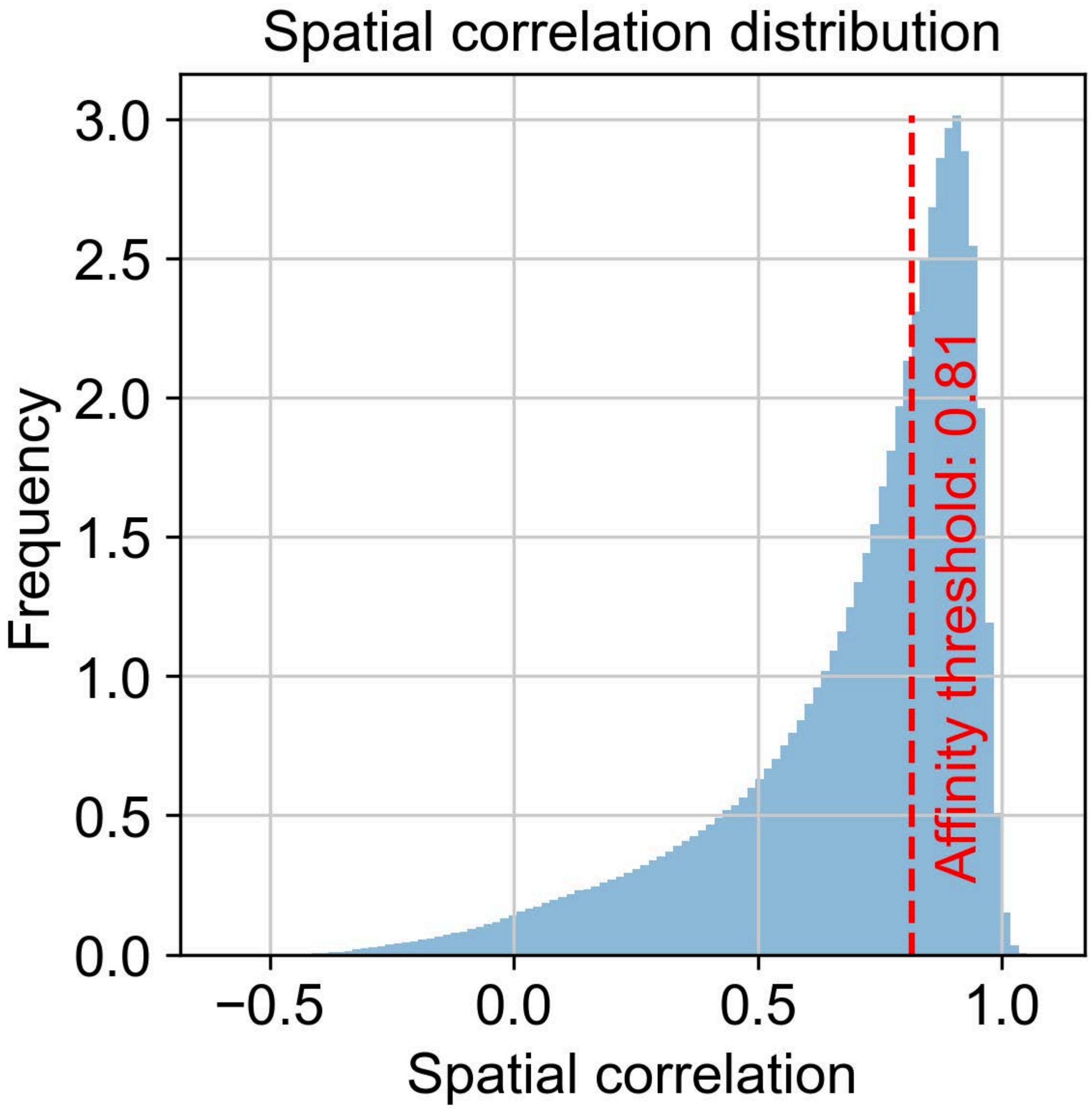


One way to visualize and analyze the SV patterns is by calculating the consensus expression using Scanpy's tl.score_genes function. This function allows you to find the related factors and similar genes for a given SV pattern. Here we visulize a spatial patterns for an example.

In [8]:

```
adata.X = adata.layers['normalized'].copy()
i = '0'
vis_list = []
vis_genes = svmodel.result_df.query('pattern == @i').sort_values('ent', ascending=F
pat_genes = svmodel.result_df.query('pattern == @i').index
sc.tl.score_genes(adata, gene_list=pat_genes, score_name = f'spatial_pat_{i}')
vis_list += [f'spatial_pat_{i}'] + vis_genes
sc.pl.spatial(
    adata, color = vis_list,
    cmap = 'RdBu_r', layer='normalized',
)
```

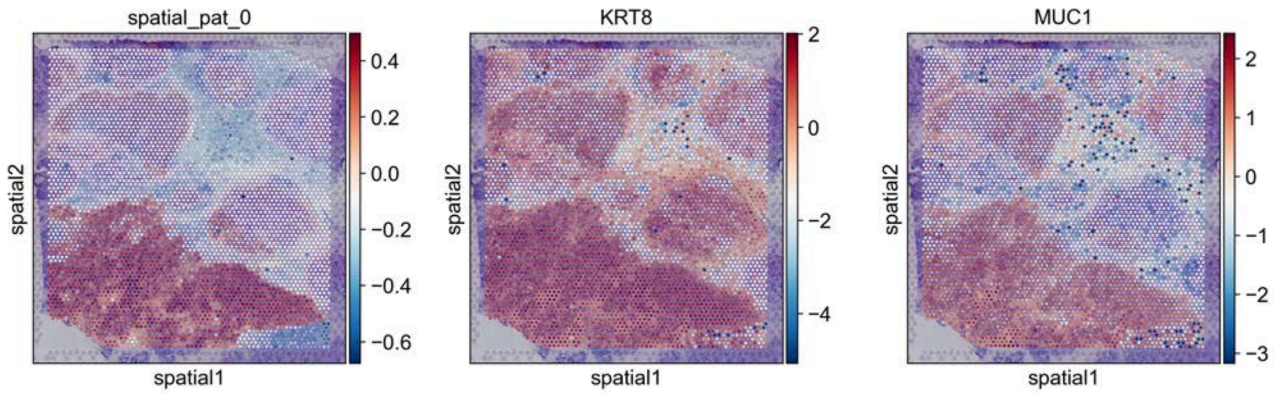


#### Utilizing Spanve aids in delineating spatial domains

Spanve considers the complex interactions between spots when detecting spatial patterns, making it a valuable tool for the crucial downstream task of spatial domain identification. In this section, we will use the imputed data from Spanve to perform clustering and identify spatial domains.

Here, we calculate the PCA embeddings for both the normalized and imputed data. The adata.var.query('spanve_spatial_features').index part ensures that we only consider the genes identified as spatially variable by Spanve when computing the PCA embeddings for the imputed data.

Next, we create a class called AutoCluster to find the best cluster number using the elbow method based on the K-means algorithm. The AutoCluster class finds the optimal number of clusters (cluster.best_k) based on the provided criteria (criteria='bic' in this case). We then use this optimal cluster number to perform K-means clustering on both the imputed and normalized PCA embeddings, storing the results in the adata.obs annotation.

Finally, we compare the results of the Spanve-aided clustering with the classical Leiden clustering approach. Here, we perform Leiden clustering on the normalized PCA embeddings, using a resolution of 0.5. We then visualize the spatial domains identified by each method (Spanve imputation with KMeans, KMeans on normalized data, and Leiden clustering) using Scanpy's sc.pl.spatial function.

By utilizing Spanve's ability to capture spatial interactions and identify spatially variable features, we can obtain more accurate spatial domain delineations (for the detailed comparison, please refer to our manuscript), which can be valuable for downstream analyses and biological interpretations.

In [9]:
```
# obtain some embeddings.
adata.obsm['normalized_pca'] = sc.pp.pca(adata.layers['normalized'])
adata.obsm['imputated_pca'] = sc.pp.pca(adata[:, adata.var.query('spanve_spatial_fe
sc.pp.neighbors(adata, use_rep='normalized_pca', metric='cosine')
sc.tl.leiden(adata, resolution=0.5, key_added='leiden_05')
```

In [10]:
```
from sklearn.decomposition import PCA
from sklearn.cluster import KMeans
from Spanve import AutoCluster

cluster = AutoCluster(init_k=3,max_k=15,criteria = 'bic')
labelx = cluster.fit_predict(adata.obsm['imputated_pca'],verbose=True)
cluster.plot_elbow()
adata.obs['imputated_KM'] = labelx.astype(str)
adata.obs['normalized_KM'] = KMeans(n_clusters = cluster.best_k).fit_predict(adata.
```

```
Sample size: 4898, using model: KMeans
finding best cluster number: 100%|██████████| 13/13 [00:02<00:00,  6.38it/s]
Best k: 7, Now predicting
```

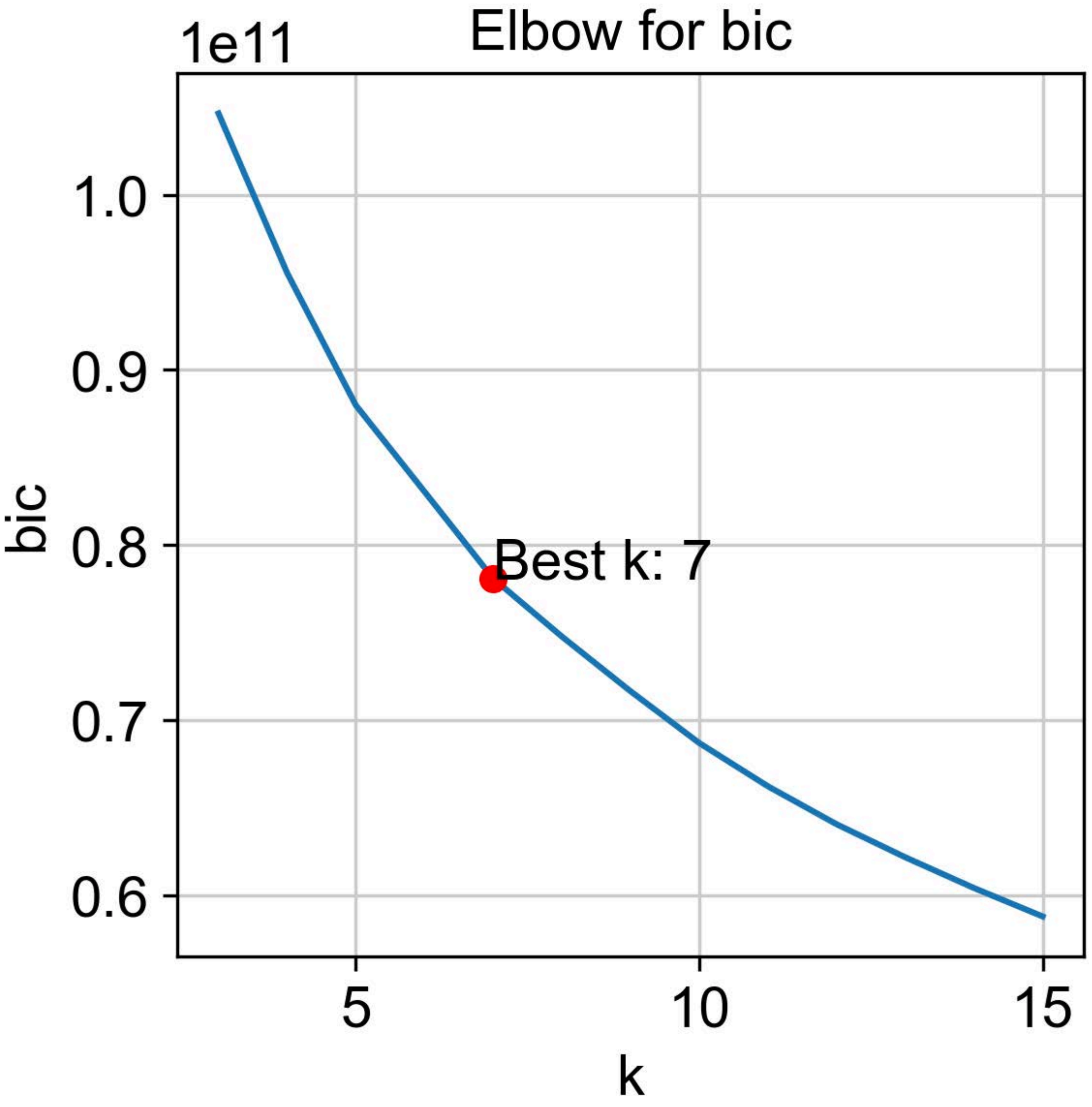


In [11]:
```python
sc.pl.spatial(
    adata, color= ['imputated_KM', 'normalized_KM', 'leiden_05'],
    legend_loc=None,
    title= ['Spanve imputation with KMeans', 'KMeans', 'Leiden'],
    ncols = 3,
)
```

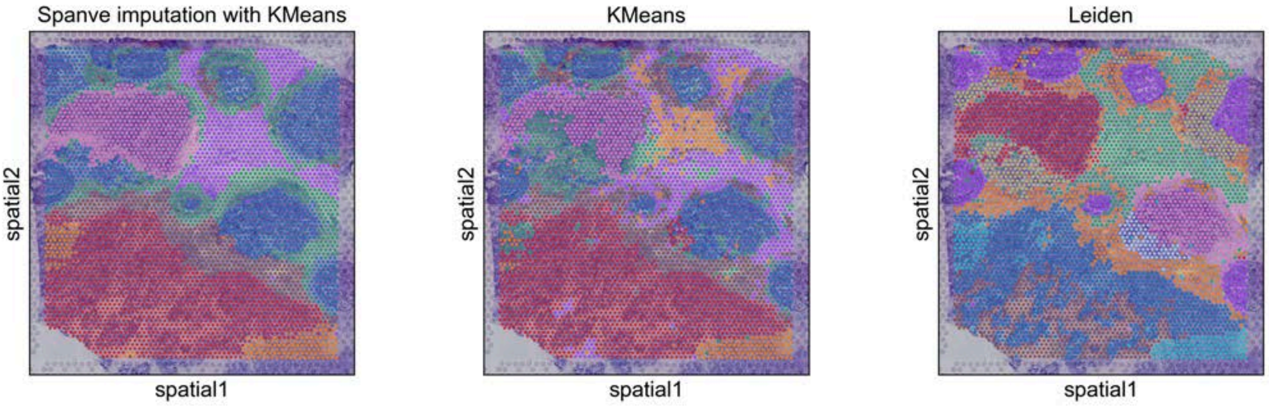


#### Spatial Colocate analysis

Spanve also provides a way to analyze the spatial co-localization patterns between gene pairs, which can indicate potential short-distance cell-cell interactions (CCI) at the spot level. This analysis complements existing studies that focus on cell communication between spot clusters.

To perform spatial colocate analysis, we first need to preprocess the data using the spatial_colocate_preprocess function provided by Spanve. This function transforms the gene expression data into a format suitable for colocate analysis, inspired by the Pearson correlation formula. Note that this step modifies the adata object.

In [12]:
```
import networkx as nx

# import commot as ct
# df_ligrec=ct.pp.ligand_receptor_database(database='CellChat', species='human', si
# df_ligrec.columns = ['ligand', 'receptor','pathway', 'type']

df_ligrec = pd.read_csv('./CellChatDB.csv')
pairs_types = []
all_search_pairs = []
for i in range(df_ligrec.shape[0]):
    ligs = df_ligrec.iloc[i,0].split('_')
    recs = df_ligrec.iloc[i,1].split('_')
    for lig in ligs:
        for rec in recs:
            if (lig not in adata.var_names) or (rec not in adata.var_names) or (lig
                continue
            all_search_pairs.append((lig, rec))
            pairs_types.append(df_ligrec.iloc[i, 3])
```

In [13]:
```
adata.X = adata.layers['counts']
scmodel = Spanve(adata.copy(), njobs = 1)
scmodel.spatial_colocate_preprocess(all_search_pairs)
scmodel.fit(verbose=True)
print("Detected significal gene pairs:",scmodel.rejects.sum())
```

```
Delete gene pair with pseduo counts < 1: -19880 pairs
Remaining: 347 pairs
#1 Expected Dist within Nodist Hypoth: 100%|██████████| 347/347 [00:00<00:00, 864.4
3it/s]
#2 Nearest Neighbors Found
#3 Computing Absolute Substract Value: 100%|██████████| 347/347 [00:00<00:00, 885.1
4it/s]
#4 Entropy Calculated
#5 G-test Performed
Write results to adata.var['spanve_spatial_features']
#--- Done, using time 1.43 sec ---#
Detected significal gene pairs: 19
```

We can visualize the colocalization patterns in a network like what we do in our manuscipt.

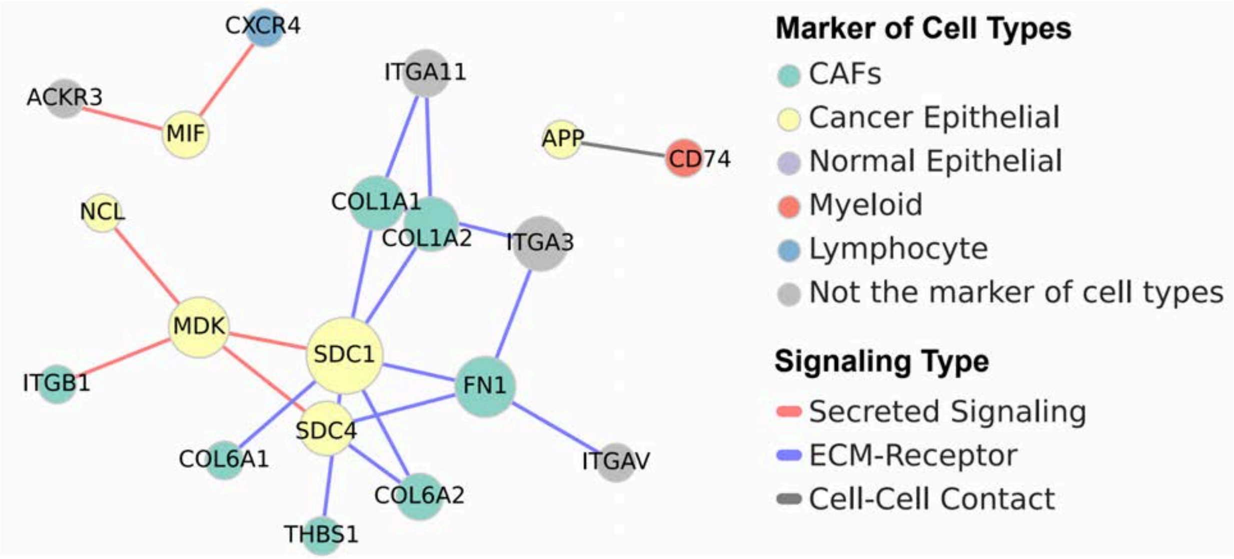


To illustrate the spatial colocalization analysis, let's consider an example with the APP and CD74 genes.

In [14]:
```
f, axes = plt.subplots(1,3, figsize = (12,4))
sc.pl.spatial(
    adata, color = 'APP', ax = axes[0],
    img_key=None, spot_size=300,
    cmap = 'Reds', show = False, colorbar_loc=None
)
sc.pl.spatial(
    adata, color = 'CD74', ax = axes[1],
    img_key=None, spot_size=300,
    cmap = 'Reds', show = False, colorbar_loc=None,
)
sc.pl.spatial(
    scmodel.adata, color = 'APP~CD74', ax = axes[2],
    spot_size=300,  cmap = 'RdBu_r', colorbar_loc=None,
    vmax = 20, vmin=-20, title = 'Co-localization of \nAPP and CD74',
    show = False
)
for ax in axes:
    ax.set_xlabel('')
    ax.set_ylabel('')
```

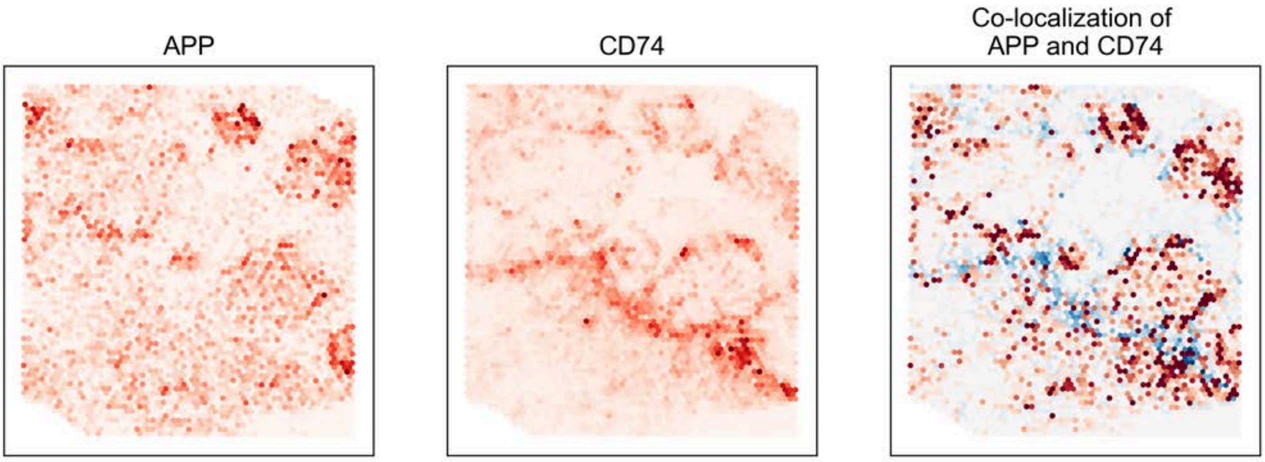
