## Supplementary file 1 for "Spanve: A Statistical Method for Detecting Downstream-Friendly Spatially Variable Genes in Large-Scale Spatial Transcriptomic Data"

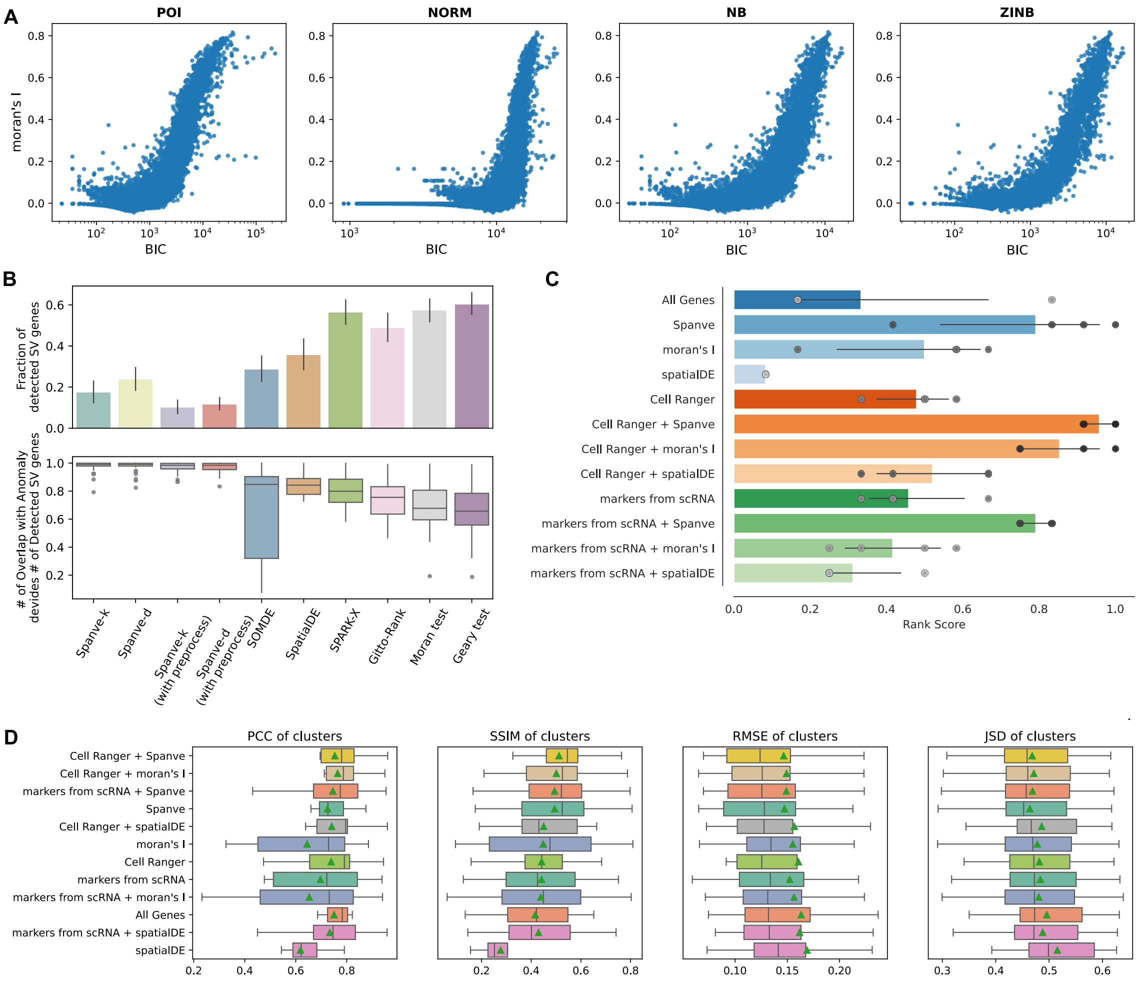


Figure S1. (A) Spatially variable (SV) genes tend to violate the assumptions of commonly used distributions for modeling gene expression in the VMOB dataset. Four distributions were fitted: Poisson (POI), Normal (NORM), Negative Binomial (NB), and Zero-Inflated Negative Binomial (ZINB). The x-axis shows the Bayesian Information Criterion (BIC), and the y-axis shows the spatial variance metric, Moran's I. (B) SV genes in 45 datasets from 10x Genomics. Upper panel: The fraction of SV genes detected by different methods across all genes in 45 datasets from 10x Genomics. Bottom panel: The fraction of overlap between SV genes and anomaly genes, defined as genes whose expression is rejected by the goodness-of-fit tests for Poisson and Normal distributions. (C) The overall ranking score and four individual metric scores for evaluating deconvolution performance on the seqFISH+ dataset using cell2location. The overall ranking score is the average of the four metric rank scores. (D) Four metrics used to evaluate deconvolution performance: Pearson Correlation Coefficient (PCC), Structural Similarity Index (SSIM), Root Mean Square Error (RMSE), and Jensen–Shannon Divergence (JSD). Higher PCC and SSIM values, and lower RMSE and JSD values, indicate better deconvolution performance.

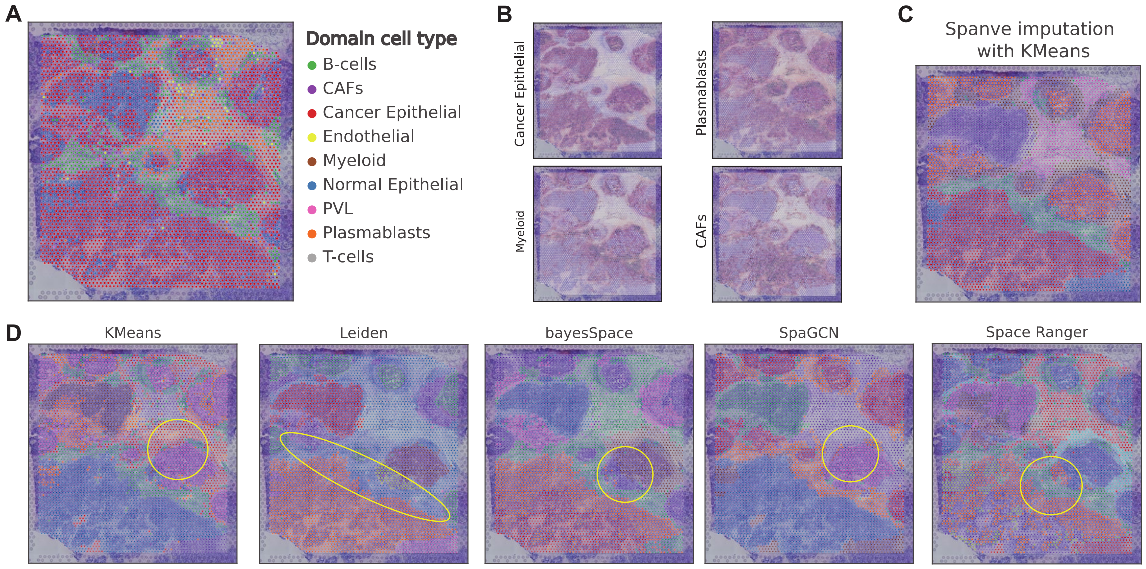


Figure S2. Comparison of human breast cancer sample clustering using different methods. (A) Domain cell types based on deconvolution results. CAFs: cancer-associated fibroblasts, PVL: perivascular-like cells. (B) Spatial distribution of key cell types aiding the identification of spatial domains. (C) Cluster result achieved from K-Means clustering on Spanve imputed data. (D) Cluster results obtained using different methods. Each result was generated using default parameters, with the cluster number set to or close to 7. The yellow circle highlights a manually annotated area that may be inaccurate. In brief, K-Means, bayesSpace, and SpaGCN may miss the layer enriched for immune cells around the tumor area; Leiden misses the layer enriched for myeloid cells; Space Ranger produces meaningless subparts of the tumor area.

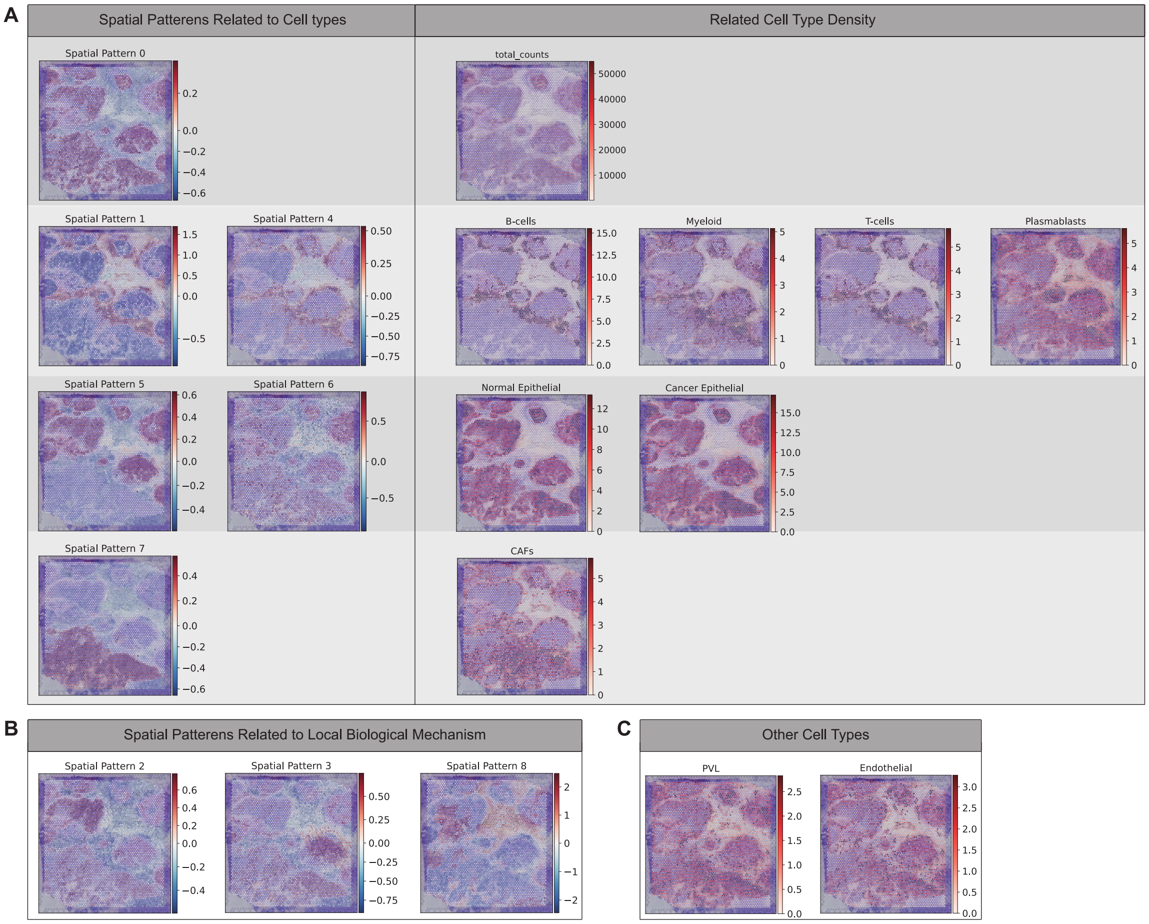


Figure S3. Spatial patterns and cell type density in human breast cancer spatial transcriptomics (ST) data. We categorized the spatial variance source into three types: cell numbers (total counts at each spot), cell type density, and local biological mechanisms. (A) The figure illustrates the scoring of spatial patterns across the tissue space, along with the related cell types and factors. (B) Spatial patterns assigned to be caused by local biogical mechanisms. (C) Other cell types that not related to the spatial patterns.

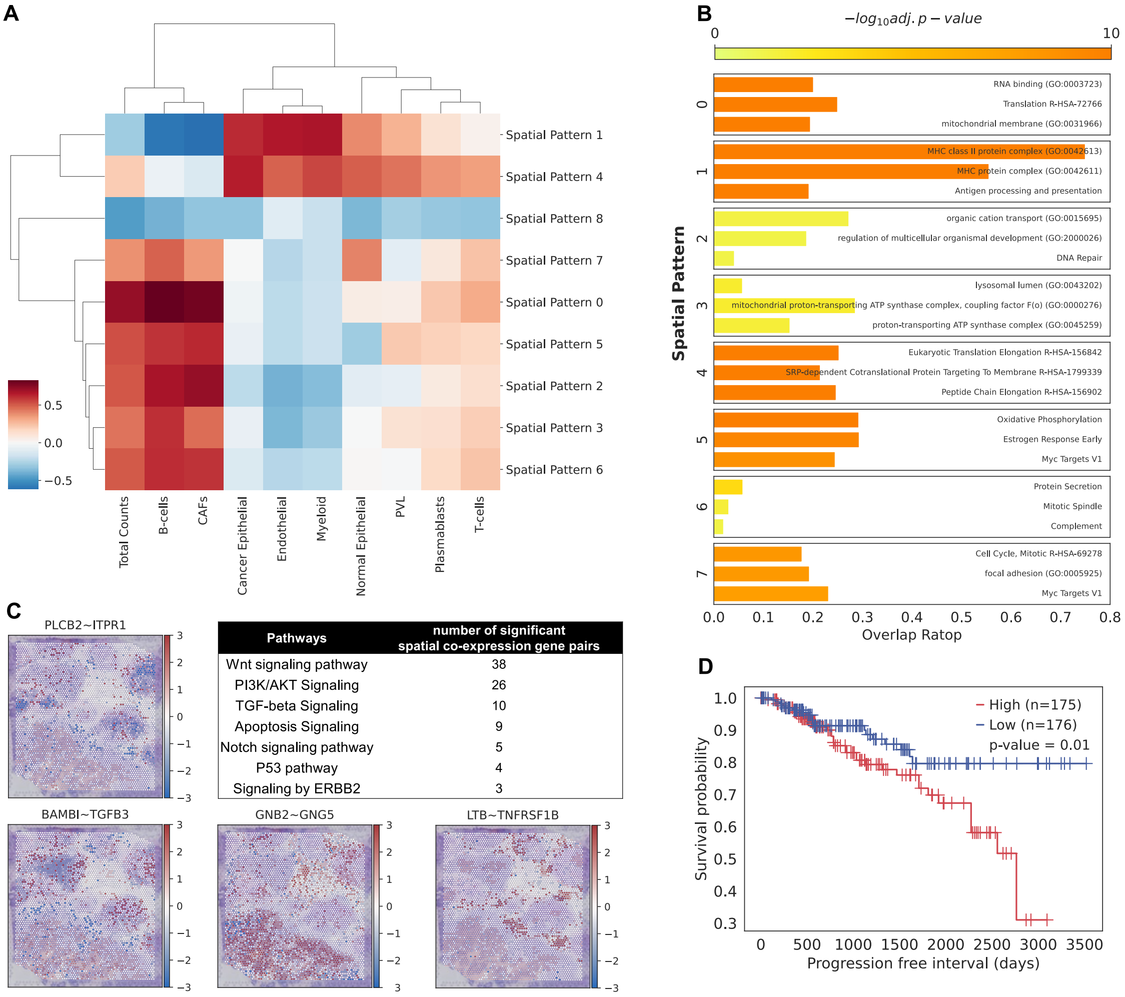


Figure S4. (A) The correlation of spatial patterns with different cell types and total counts. (B) Gene set enrichment analysis for each spatial pattern. Three most significant items are listed for each pattern. Notably, spatial pattern 8 did not yield any significant items. The gene annotations include Gene Ontology, KEGG pathway, Reactome, and MSigDB Hallmark. (C) Spatial co-localization in human breast cancer spatial transcriptomics (ST) data. The table shows the number of gene pairs with significant co-localization in the sample identified by Spanve for the seven most studied pathways. The other spatial scatter plots display examples of typical spatially co-expressed genes belonging to these 7 pathways, where red spots indicate positive co-expression and blue spots indicate negative co-expression. The x- and y-axes represent spatial coordinates of the spots, and the spot color shows the co-localization level of the genes. (D) Relationship between the signature Cluster 4 and survival in TCGA-BRCA data. The p-value is obtained from the log-rank test. The high- and low-expressed groups are determined by the upper and lower quantiles of the average scaled expression, respectively.
